# Maternal obesity disrupts glucose oxidation and mitochondrial fission in the myometrium at term gestation

**DOI:** 10.64898/2026.11.16.766658

**Authors:** Kevin K Prifti, Jordan L Rich, Amelia M Giecek, Ruth M Gill, Terri A Pietka, Wandy L Beatty, Antonina I Frolova

## Abstract

Women with obesity experience delayed spontaneous labor onset, slow labor progression, and elevated risk of labor dystocia, but the underlying mechanisms are poorly understood. We previously established a mouse model of diet-induced obesity that exhibits decreased uterine contractility, labor dystocia, and elevated fatty acids in the myometrium. Here, we used this model to determine how maternal obesity affects energy production and mitochondrial homeostasis in the myometrium at term. Myometrium tissue and myometrial smooth muscle cells from diet-induced obese mice at term showed decreased pyruvate utilization, inhibition of glucose oxidation, and disruption of mitochondrial fission and mitophagy. Treatment with medium-chain fatty acids in both murine and human primary myometrial cells resulted in significant decreases in abundance of mitochondrial fission proteins. Finally, mitochondrial spare capacity in human myometrial cells decreased with body mass index of the patient providing the cells. Taken together, these findings indicate that fatty acid accumulation dysregulates energy metabolism and mitochondrial homeostasis in the myometrium and may contribute to dysfunctional labor in women with obesity.

## INTRODUCTION

In the U.S., over one-third of childbearing women have obesity (body mass index [BMI] ≥ 30 kg/m^2^)^1^. Compared to women without obesity, these patients have lower rates of spontaneous labor at term, resulting in higher rates of induction and prolonged gestation, which increases the risk of neonatal complications^2–5^. They also tend to have slower labor progression and longer labor durations, leading to higher rates of cesarean delivery^6–9^. Several pieces of evidence suggest that maternal obesity results in dysfunctional contractility of the uterine smooth muscle (myometrium) during labor. First, despite higher rates of use of uterotonics to augment labor, maternal obesity is associated with lower rates of vaginal delivery due to failure of labor induction ^10–12^. Second, women with obesity have elevated rates of postpartum hemorrhage^13^, suggesting poor ability of the uterus to contract and prevent bleeding. Third, myometrial tissues obtained from women with obesity undergoing cesarean delivery have less contractile force and frequency than myometrium from women without obesity^14^.

Dysfunctional myometrial contractility in obesity could reflect altered nutrient availability, as the abundance of fatty acids is generally highly elevated, and inability to generate sufficient energy to achieve the contractility required to dilate the cervix and expel the fetus. Energy generation in all cells relies on mitochondrial oxidative phosphorylation, which most commonly uses glucose or fatty acids as the substrates. However, myometrial cells from non-obese women rely primarily on glycolysis and glucose oxidation for energy production in both quiescent and oxytocin-induced contractile states and are limited in their ability to use fatty acids^15^. Dysfunctional myometrial contractility in obesity could also reflect impaired mitochondrial function. Mitochondrial homeostasis is maintained by degradation of damaged mitochondria via mitophagy, as well as the balance of fusion and fission, which regulate mitochondrial size, shape, and number in response to local demand for ATP^16,17^. Obesity is associated with disruption of mitochondrial dynamics in skeletal muscle, and exposure to a variety of lipids results in mitochondrial fragmentation in several cell types^18–20^. Excess aggregation of lipids in skeletal and cardiac muscle also results in increased fatty acid beta-oxidation and inhibition of glucose breakdown and oxidation. The effects of obesity on mitochondrial function and energy production in the myometrium is unknown.

Here, we used a mouse model and human samples to fill this gap. First, we used a mouse model of maternal diet-induced obesity that exhibits decreased uterine contractility, labor dystocia, and higher fatty acid content in the myometrium^21^. We assessed metabolic activity in murine myometrial fibers and primary myometrial cells. Additionally, we quantitatively assessed mitochondrial morphology and ultrastructure in murine myometrial tissue and quantitated abundance of proteins associated with mitochondrial fusion, fission, and mitophagy. We then quantitated mitochondrial respiration and mitochondrial dynamics in primary human myometrial cells exposed to exogenous fatty acids. Finally, we quantitated mitochondrial respiration differences in primary human myometrial cells from women with and without obesity. Together, our data indicate that maternal obesity disrupts glucose utilization and mitochondrial morphology, ultrastructure, and DRP1-regulated fission in the myometrium and that this is, at least in part, due to exposure to excess fatty acids.

## MATERIALS AND METHODS

### Animals and diet

All animal procedures were approved by the Institutional Animal Care and Use Committee at Washington University School of Medicine (protocol number 20-0339) and conformed to NIH guidelines. Mice were housed in ventilated cages under controlled temperature and humidity and on a 12-h light (06:00-18:00):12-h dark cycle (18:00-06:00). Four-week-old female C57BL/6J mice (The Jackson Laboratory, Bar Harbor, ME) were fed either control chow (CON; TestDiet® Formula 58R0; 11% fat, 73% carbohydrates [0.3% sucrose], and 15% protein) or an obesogenic diet (DIO; TestDiet® Formula 58R3: 59% fat, 26% carbohydrates [17% sucrose], and 15% protein) for 10 weeks. All mice that consumed the obesogenic diet were included in the DIO experimental group, irrespective of the amount of weight gained. Females in both dietary groups were then mated to control-chow-fed males overnight. Pregnancy was confirmed by identification of semen plug at 08:00 and was recorded as pregnancy day 0.5 (P0.5). During mating, all mice were fed control chow. After mating, the females were fed their respective diets throughout pregnancy.

### Isolation and cell culture of primary mouse myometrial smooth muscle cells

Uteri were collected on P18.5 (term non-labor) at 09:00-11:00. The implantation sites closest to the cervix were designated as C-0, the next as C-1, etc. As described^21^, for each implantation site, fetal tissue, amnion, and placenta were removed, and the remaining uterine tissue was flash frozen in liquid nitrogen and stored until further analysis. C-1 sites were used for primary mouse myometrial smooth muscle cell (mMSMCs) isolation. The uterine tissue was washed in ice-cold sterile Hanks’ Balanced Salt Solution (Gibco) supplemented with 5 μg/mL amphotericin B (Corning) and 50 μg/mL gentamicin. Tissue was cut into 1-2 mm pieces and incubated under constant rotation at 37 °C for 60 min in digestion solution: 1 mg/mL collagenase 1A and XI (Sigma) with 50 μg/mL gentamicin diluted in 4 mL DMEM with 1% charcoal stripped fetal bovine serum (FBS) (Gibco). The solution was then filtered through a 100 μM sterile cell strainer (CellTreat Scientific Products). Collagenase activity was stopped by adding 20 mL of warm 10% FBS + DMEM through the cell strainer. The cell suspension was centrifuged for 5 minutes and the supernatant was discarded. The cell pellet was resuspended in 3 ml of ACK Lysis Buffer (Gibco). After 5 minutes, 3 mL of 10% FCS + DMEM was added, the cell suspension was centrifuged again, and the supernatant discarded. Cells were plated and cultured in DMEM supplemented with smooth muscle cell growth media 2 mix (Promo Cell) and 50 μg/mL gentamicin. All cells were frozen at P0 and thawed to use as needed.

### Isolation and culture of primary human myometrial smooth muscle cells

The Washington University in St. Louis Human Research Protection Office approved this study (IRB #202511061). Deidentified human myometrial tissue was procured under a waiver of consent from the Washington University Reproductive Specimen Processing and Banking (ReProBank) core. Primary human myometrial smooth muscle cells (hMSMCs) were isolated from biopsies collected from the upper aspect of the hysterotomy performed at the time of term scheduled (unlabored) cesarean delivery as previously described^15^. All cells were frozen at P1 and thawed to use as needed between P2 and P4.

### In vitro fatty acid treatments

Primary mMSMCs or hMSMCs were plated in 6-well culture plates and allowed to adhere overnight. Then, cells received a single dose of either vehicle (DMSO, at final concentration 0.32%) or 250 µM octanoic acid (OA, C8:0, Cayman Chemical), were incubated for 18 hours, and were lysed and used for protein analysis.

### High-resolution respirometry of myometrial fibers

An Oroboros Oxygraph (Oroboros Instruments) was prepared as previously described^22^. Mouse C-0 implantation sites collected on P18.5 (term non-labor) were dissected, finely chopped, collected in BIOPS solution (10 mM Ca-EGTA buffer, 20 mM imidazole, 20 mM taurine, 50 mM K-MES, 0.5 mM DTT, 6.56 mM MgCl_2_, 5.77 mM ATP, 15mM phosphocreatine), permeabilized with 5 mg/ml saponin for 20 min, and rinsed with MiR05 buffer for 15 min. Tissue was the blotted dry and weighed, and 15-20 mg was placed in an Oxygraph chamber with 2 ml MiR05 buffer supplemented with 20 mM creatine monohydrate. Before the chamber was fully sealed, the tissue was oxygenated with 1 ml of 100% oxygen. After oxygen levels equilibrated, the oxygen consumption rate (OCR) was measured as a readout for substrate utilization in the mitochondria and normalized to tissue weight. Myometrial fibers were exposed to either carbohydrate (pyruvate) or fatty acid (octanoyl-carnitine) substrates, and OCR was recorded by sequential injection of the following: glutamate/malate (complex I stimulator), ADP (converts to ATP, activates oxidative phosphorylation [OXPHOS]), succinate (complex II stimulator), cytochrome C (to assess potential mitochondrial damage), and FCCP (uncoupling agent). The following parameters were analyzed: 1) Complex I State 2 respiration (LEAK) was calculated after the addition of malate/glutamate/pyruvate (aerobic glycolytic protocol) or malate/glutamate/octanoyl-carnitine (fatty acid protocol). 2) Complex I State 3 respiration (CI) was calculated after the addition of ADP. 3) Complex I + II State 3 respiration (OXPHOS) was calculated after the addition of succinate. 4) Complex I + II + Cytochrome State 3 (CYT C) respiration was calculated after the addition of Cytochrome C. 5) Maximal Respiratory Rate (MAX ETS) was calculated after FCCP titration at the highest O_2_ flux rate. 6) Respiratory Control Ratio (RCR) was calculated as OXPHOS:LEAK ratio. Any samples demonstrating an increase in O_2_ flux rate of 15% or higher after Cytochrome C injection were excluded from analysis due to compromised outer mitochondrial membrane^22^.

### Preparation of cells for bioenergetic analysis

To prepare Mitochondrial Stress Test and Substrate Oxidation Test medium, Seahorse XF DMEM medium, pH 7.4 (Agilent), was supplemented with 1 mM sodium pyruvate (Corning), 2 mM glutamine (Corning), and 10 mM glucose (Sigma). To prepare Glycolysis Stress Test medium, Seahorse XF DMEM medium, pH 7.4 (Agilent), was supplemented with 2 mM glutamine (Corning). At passages 2–4, 30,000 mMSMCs were plated in Seahorse 96-well culture plates (Agilent). After 18 hours, cell culture growth medium was removed, and cells were washed twice with pre-warmed assay medium to remove bicarbonate. After washing, cells were placed in 180 μL of assay medium, and the cell culture plate was placed in a 37 °C non-CO_2_ incubator for 45–60 min until loading it into the Seahorse XFe96 Analyzer (Agilent).

### Extracellular flux analysis

One day before the experiment, a Seahorse XFe96 sensor cartridge (Agilent) was hydrated by filling each well with 200 uL of calibration solution (Agilent), sensors were submerged into it, and it was placed in a non-CO_2_ incubator at 37 °C overnight. On the day of the experiment, the sensor cartridge was taken out of the non-CO_2_ incubator, and the injection ports were filled with appropriate compounds for each assay. For the Mitochondrial Stress Test, the following compounds were used: oligomycin (OG, ATP synthase complex V inhibitor; Cayman Chemical), trifluoromethoxy carbonyl cyanide phenylhydrazone (FCCP, uncoupling agent that abolishes the proton gradient), and a combination of rotenone (ROT, complex I inhibitor; Sigma) with antimycin A (AA, complex III inhibitor; Cayman Chemical) to interrupt mitochondrial respiration. For the Glycolysis Stress Test, the following compounds were used: glucose (saturating dose fueling glycolysis; Sigma), oligomycin (mitochondrial ATP synthase complex V inhibitor, which increases reliance on glycolysis), 2-deoxy-glucose (2-DG, competitive inhibitor of glucose, which inhibits glycolysis; Sigma). Compounds were reconstituted, aliquoted, and store at -80 °C. On the day of the assay, an aliquot was thawed and used to prepare the stock solutions in assay medium and then further diluted to 10X concentrations before being loaded into the ports. The following final concentrations were used: 1.5 μM OG, 0.5 μM ROT, 0.5 μM AA, 10 mM glucose, and 50 mM 2-DG. FCCP titration was performed as described^15^, and optimal concentrations for mMSMCs and hMSMCs were found to be 1.0 μM and 4.0 μM, respectively. The loading volumes of each 10X compound into the injection ports were 20 μL for Port A, 22 μL for Port B, 25 μL for Port C, and 27 μL for Port D. When calibration was complete (after about 20 min), the utility plate was replaced with the culture plate. After a short mixing period, the first set of measurements were recorded under basal conditions. Three measurements 8 min apart of oxygen consumption rate (OCR) and extracellular acidification rate (ECAR) were recorded between each injection.

### Calculation of metabolic parameters for mitochondrial function

Mitochondrial stress test analysis provided direct measurements of basal respiration, ATP-linked respiration, maximal respiration, spare capacity, non-mitochondrial respiration, and proton leak. Averages of individual well results (n = 3) for each sample were calculated based on OCR measurements. Because equal numbers of cells were plated per well for each sample, Seahorse data were internally normalized to cell count. ECAR is driven by lactate accumulation in the extracellular medium. For the mitochondrial stress test assay, a proportional correlation between ECAR and lactate production was assumed, and changes in ECAR were used to indirectly reflect glycolytic capacity. Estimations of energy generation via oxidative phosphorylation vs. glycolysis were calculated at the basal rate of acidification (OCR/ECAR at baseline), after adding oligomycin to inhibit ATP-linked respiration (OCR/ECAR after OG), and after adding FCCP to induce maximal ATP requirement (OCR/ECAR at max). Area-under-the-curve (AUC) was calculated as the total mitochondrial respiration OCR above baseline, which was set as the mean of values recorded upon addition of ROT/AA.

### Calculation of metabolic parameters for glycolytic function

The glycolysis stress test provided direct measurements of basal glycolysis, glycolytic reserve, glycolytic capacity, and non-glycolytic acidification. Averages of individual well results (n = 3) for each sample were calculated. Because equal numbers of cells were plated per well for each sample, Seahorse data were internally normalized to cell count.

### Western blot analyses

Mouse C-1 implantation sites were used for protein analysis. Tissue was homogenized with a Bullet Blender 24 (Next Advance Inc.) at 4°C in 250 μl lysis buffer (50 mM Tris HCl [pH 8.0], 150 mM NaCl, 1 mM EDTA, 0.5% Nonidet P-40, 2% Glycerol, complete Protease Inhibitor Cocktail [Roche Applied Science], and Phosphatase Inhibitor Mini Tablets [ThermoFisher]) per 5 mg of tissue. Cell lysates were collected from 6-well culture plates using 200 μl lysis buffer per well. Lysates were centrifuged at 14,000 g for 10 min at 4°C. Supernatants were stored at -80°C. Protein concentration was quantified with the BCA Protein Assay Kit (Pierce). Proteins were then separated by SDS-PAGE in 4% to 12% Bolt Bis-Tris gels (ThermoFisher) and transferred to PVDF. Membranes were stained for total protein with REVERT™ Kit (LI-COR) according to the manufacturer’s instructions. Briefly, membranes were incubated in REVERT for 5 minutes, washed twice for 30 seconds, dried for 1 minute at room temperature (RT) and imaged immediately. Membranes were then rehydrated in 1x Tris-buffered saline (TBS) and blocked in 5% nonfat dry milk powder in TBS for 1 h. Blots were incubated in 5% nonfat dry milk in TBS with 0.01% Tween 20 overnight at 4 °C with primary antibodies and then incubated for 1 h at RT with appropriate secondary antibodies. Membranes were stained with antibodies against GLUT1 (1:2000, Abcam Cat# ab115730, RRID:AB_10903230), GLUT3 (1:1000, Abcam Cat# ab191071, RRID:AB_2736916), GLUT4 (1:2000, Proteintech Cat# 66846-1-Ig, RRID:AB_2882186), HK1 (1:1000, Proteintech Cat# 19662-1-AP, RRID:AB_10859778), PKM2 (1:1000, Proteintech Cat# 15822-1-AP, RRID:AB_1851537), LDHA (1:1000, Proteintech Cat# 19987-1-AP, RRID:AB_10646429), MPC1 (1:1000, Cell Signaling Technology Cat# 14462, RRID:AB_2773729), MPC2 (1:1000, Cell Signaling Technology Cat# 46141, RRID:AB_2799295), Total PDH (1:1000, Cell Signaling Technology Cat# 3205, RRID:AB_2162926), Phosphorylated PDH (Ser232) (1:1000, Sigma-Aldrich Cat# AP1063, RRID:AB_10616070), Phosphorylated PDH (Ser300) (1:1000, Millipore Cat# AP1064, RRID:AB_10618090), Phosphorylated PDH (Ser293) (1:1000, Millipore Cat# ABS204, RRID:AB_11213668), BNIP3 (1:1000, Proteintech Cat# 68091-1-Ig, RRID:AB_2918828), LC3B-I/II (1:1000, (Cell Signaling Technology Cat# 2775, RRID:AB_915950)), FUNDC1 (1:1000, Proteintech Cat# 28519-1-AP, RRID:AB_2935469), PINK1 (1:1000, Cell Signaling Technology Cat# 6946, RRID:AB_11179069), TOM20 (1:1000, Cell Signaling Technology Cat# 42406, RRID:AB_2687663), MFN1 (1:1000, Cell Signaling Technology Cat# 14739, RRID:AB_2744531), MFN2 (1:1000, Cell Signaling Technology Cat# 11925, RRID:AB_2750893), OPA1 (1:1000, Cell Signaling Technology Cat# 80471, RRID:AB_2734117), MFF (1:1000, Cell Signaling Technology Cat# 84580, RRID:AB_2728769), DRP1 (1:1000, Cell Signaling Technology Cat# 8570, RRID:AB_10950498), FIS1 (1:1000, Cell Signaling Technology Cat# 32525, RRID:AB_2927422), followed by anti-mouse IgG (H+L) (Jackson ImmunoResearch Labs Cat# 115-035-003, RRID:AB_10015289) or anti-rabbit IgG (H+L) (Jackson ImmunoResearch Labs Cat# 111-035-003, RRID:AB_2313567) HRP-conjugated secondary antibody. The signal was detected with ECL Western Blot Substrate (Thermo Fisher), membranes were scanned on a ChemiDoc MP (Bio-Rad Laboratories), and signals were quantitated with Image Lab (Bio-Rad Laboratories). Values for *GLUT1, GLUT3, GLUT4, HK1, PKM2, LDHA, MPC1, MPC2, BNIP3, LC3B-I/II, FUNDC1, PINK1, TOM20, MFN1, MFN2, OPA1, MFF, DRP1, FIS1* proteins were normalized to total protein. Phosphorylated PDH was normalized to total PDH. In two cases, membranes were cut and probed with two antibodies, so intensities were normalized to the same total protein blot: 1) HK1 (Figure 4D) and BNIP3 (Figure 6B); 2) PKM2, LDHA (Figure 4D) and FUNDC1 (Figure 6B).

### Transmission electron microscopy

Mouse C-2 implantation sites were fixed in 2% paraformaldehyde/2.5% glutaraldehyde (Ted Pella Inc.) in 100 mM cacodylate buffer, pH 7.2 for 2 hr at room temperature and then overnight at 4°C. Samples were washed in cacodylate buffer and postfixed in 1% osmium tetroxide (Ted Pella Inc.)/ 1.5% potassium ferricyanide (Sigma) for 1 hr. Samples were then rinsed extensively in dH20 before *en bloc* staining with 1% aqueous uranyl acetate (Ted Pella Inc.) for 1 hr. Following several rinses in dH20, samples were dehydrated in a graded series of ethanol and embedded in Eponate 12 resin (Ted Pella Inc.). Sections (95 nm) were cut with a Leica Ultracut UCT ultramicrotome (Leica Microsystems Inc.), stained with uranyl acetate and lead citrate, and viewed on a JEOL 1200 EX transmission electron microscope (JEOL USA Inc.) equipped with an AMT 8 megapixel digital camera and AMT Image Capture Engine V602 software (Advanced Microscopy Techniques). Mitochondrial morphology and ultrastructure were qualitatively and quantitatively assessed in ImageJ^23,24^. To prevent experimental bias, images were coded, and a masked investigator quantitated the following parameters: mitochondrial length, width, area, perimeter, shape, circularity index, roundness index, integrated density, distance from the plasma membrane, and cristate quality. Additionally, mitochondria-endoplasmic reticulum (ER) contact length, ER quality, and percent mitochondria associated with ER were assessed.

### Statistical analysis

Results were averaged for each treatment group. Data are presented as means ± standard errors of the means (SEM). Significance was determined by student t-test or one-way ANOVA, with Holm-Sidak multiple comparison test in Prism 11 software (GraphPad) as appropriate. Linear regression was performed and 95% confidence intervals for the predicted means were estimated using Stata Version 18 (StataCorp). Significant differences compared to controls are annotated as follows: *P < 0.05, **P < 0.01, ***P < 0.001, and ****P < 0.0001.

### Study approval

Washington University in St. Louis Human Research Protection Office approved this study (IRB #202511061). Deidentified human myometrial samples were obtained under a waiver of consent from the Reproductive Specimen Processing and Banking core (ReProBank) at Washington University School of Medicine.

### Data availability

The authors agree to share all publication-related data.

## RESULTS

### Myometrial fibers from obese mice have reduced mitochondrial respiration in the presence of pyruvate

To determine the impact of obesity on myometrial metabolism, we used an established model^21^ in which we fed four-week-old female mice a control (CON) diet or a diet-induced obesity (DIO) diet for 10 weeks. We then mated them with CON-fed males and maintained them on their respective diets during pregnancy. At gestational day 18.5 (term but before labor initiation), we isolated myometrial tissue from CON and DIO mice and analyzed mitochondrial respiration using an Oroboros Oxygraph (Figure 1A). To assess the abilities of myometrial fibers to utilize carbohydrates and fatty acids, we provided them with either the pyruvate or octanoyl carnitine and measured oxygen consumption rate (OCR) following a series of mitochondrial complex stimuli (Figure 1B–C). After complex I stimulation and ADP injection, OCR did not differ between CON and DIO myometrial fibers in either substrate group (Figure 1D–E). After stimulation of oxidative phosphorylation, both CON and DIO myometrial fibers were equally capable of utilizing octanoyl carnitine. However, DIO myometrial fibers exhibited significantly lower OXPHOS activity in the presence of pyruvate than CON fibers (Figure 1F). Furthermore, DIO fibers exhibited significantly lower OCR at maximal respiration in the presence of pyruvate than in the presence of octanoyl carnitine (Figure 1G). We observed no differences in the respiratory control ratio (RCR) between CON and DIO myometrial fibers in either substrate group (Figure 1H). Taken together, these data indicate that myometrial fibers from DIO mice have normal global mitochondrial substrate oxidation and ATP coupling but lower carbohydrate oxidation than fibers from CON mice.

**Figure 1:**
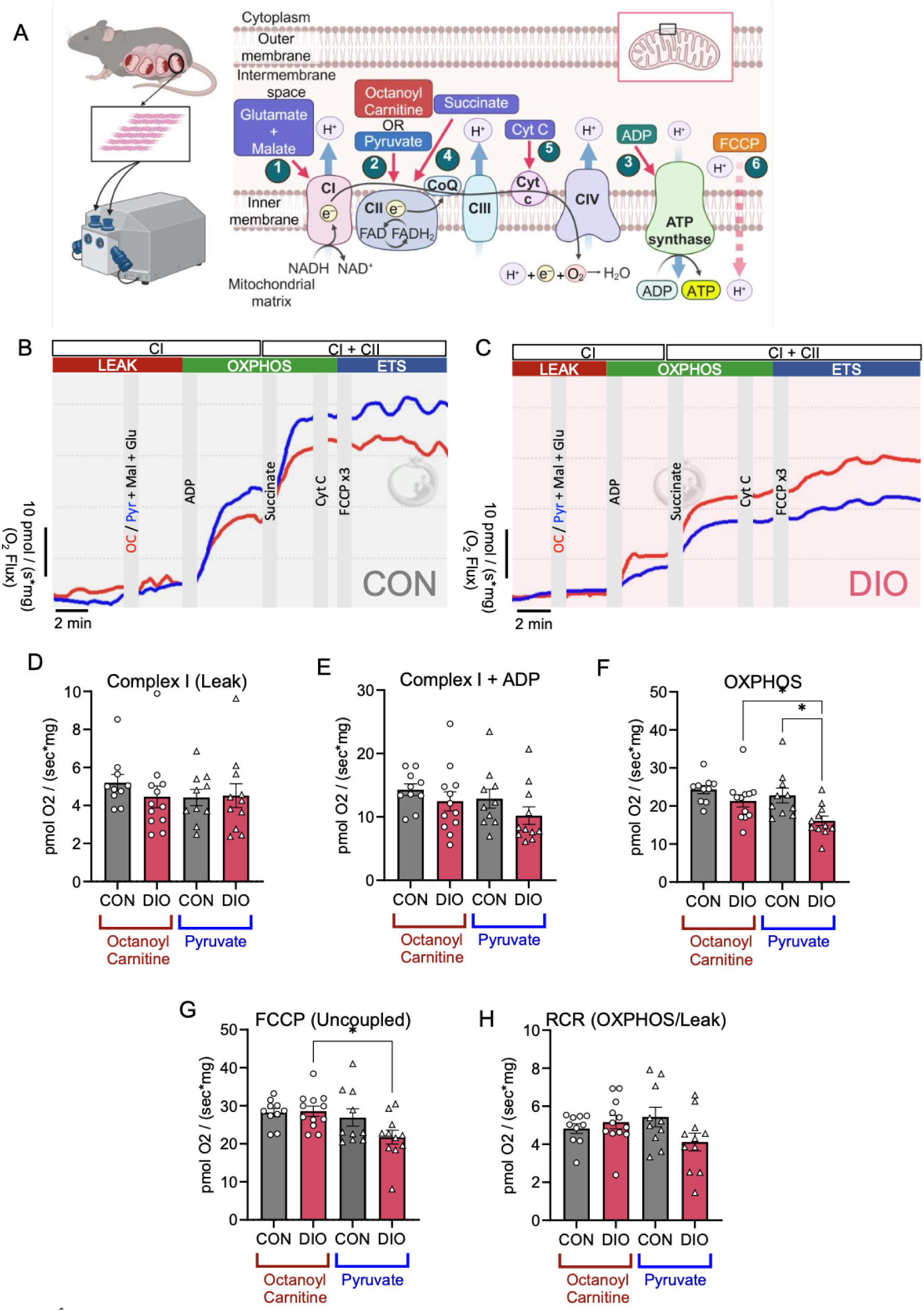
Mitochondrial respiration is decreased in DIO myometrial fibers in the presence of pyruvate. (A) Experimental schematic for high-resolution respirometry with Oroboros O2K in myometrial fibers. Circled numbers indicate drug injection order. Representative *ex vivo* myometrial fiber respirometry traces from (B) CON (n=10, grey), (C) DIO (n=11, pink) treated with pyruvate (Pyr, blue) or octanoyl carnitine (OC, red). Quantification of oxygen flux of CON + octanoyl carnitine (OC, n=10, black), DIO + OC (n=12, pink), CON + pyruvate (n=10, grey), DIO + pyruvate (n=11, pink), (D) complex I (LEAK), (E) Complex I + ADP, (F) oxidative phosphorylation (OXPHOS), (G) trifluoromethoxy carbonyl cyanide phenylhydrazone (FCCP, uncoupled), and (H) respiratory control ratio (RCR, OXPHOS/LEAK). Results are presented as means +/- SEM. \**P* < 0.05 by one-way ANOVA with Holm-Sidak multiple comparison test. ADP, adenosine diphosphate; ATP, adenosine triphosphate; CI, complex I; CII, complex II; Cyt C, cytochrome c; ETS, electron transport system; Glu, glutamate; Mal, malate; Succ, succinate.

### Myometrial cells from obese mice have reduced mitochondrial energy output

In addition to myometrial cells, the uterus contains stromal and immune cells^25^. To examine metabolic differences specifically in myometrial cells, we isolated primary murine myometrial smooth muscle cells (mMSMCs) from CON and DIO mice at term gestation and used the Seahorse XF Cell Mito Stress Test to measure mitochondrial OCR and associated bioenergetic parameters (Figure 2A). Whereas mMSMCs from CON uteri showed a typical OCR curve, mMSMCs from DIO uteri had a blunted OCR curve (Figure 2B) with a significantly lower area under the curve (Figure 2C). Basal respiration (Figure 2D), ATP-linked respiration (Figure 2E), maximal respiration (Figure 2F), spare capacity (Figure 2G), non-mitochondrial respiration (Figure 2H), and proton leak (Figure 2I) were all significantly lower in mMSMCs from DIO mice than in cells from CON mice, indicating that mMSMCs from DIO mice have lower baseline energy production and lower mitochondrial bioenergetic capacity than cells from CON mice. We observed no difference in the coupling efficiency between CON and DIO myometrial cells (Figure 2J), suggesting that the overall mitochondrial efficiency was not different between groups. Additionally, the OCR to extracellular acidification rate (ECAR) ratios (Supplementary Figure S1A–D) did not differ between CON and DIO groups at basal respiration or after oligomycin addition, but were significantly lower at maximal respiration in mMSMCs from DIO mice compared to CON, suggesting that they have a lower rate of oxidative phosphorylation or higher reliance on glycolysis at maximal respiratory demand.

**Figure 2:**
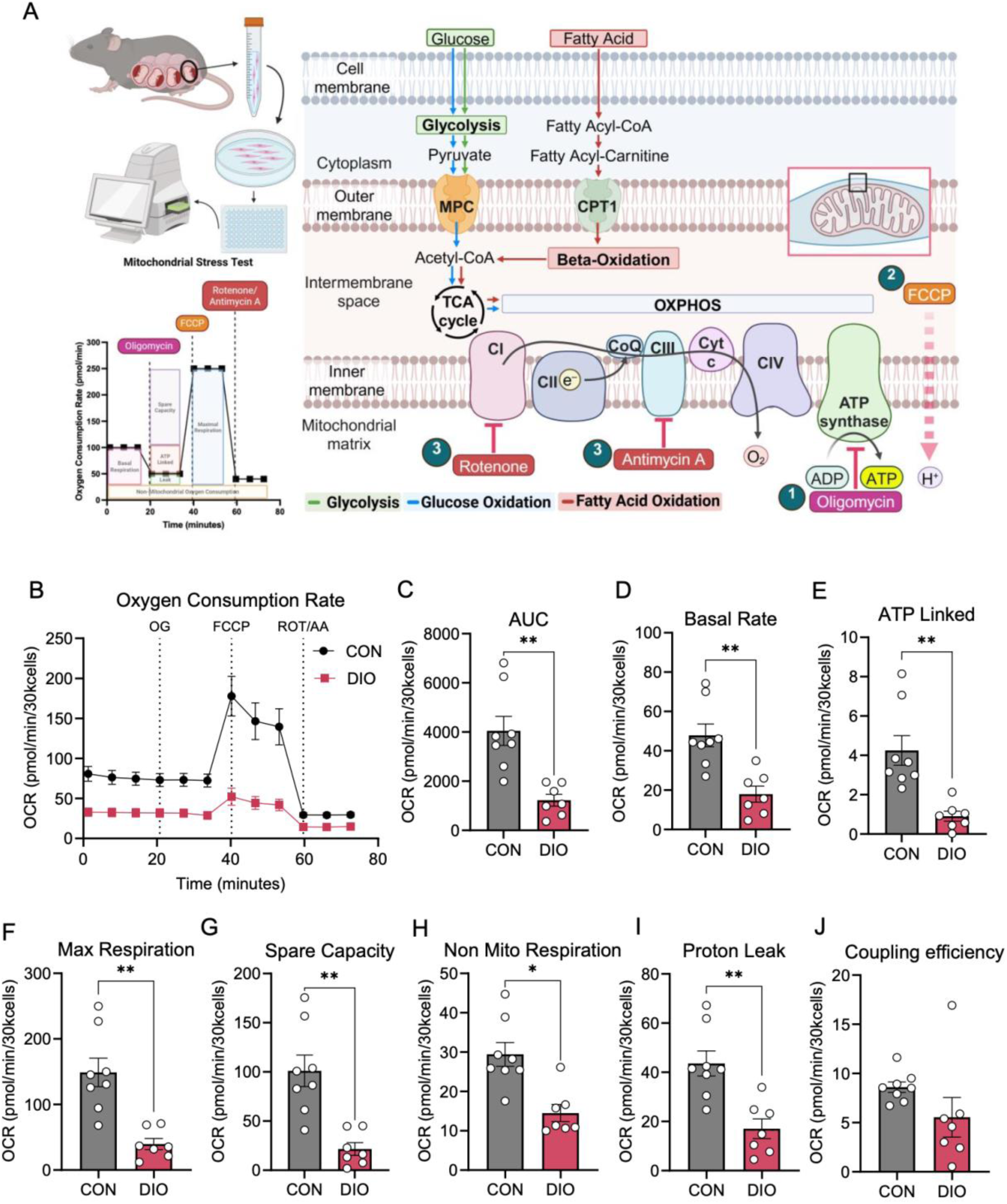
Mitochondrial bioenergetic parameters are decreased in DIO myometrial cells. (A) Left, experimental schematic for mitochondrial respiration with Agilent Seahorse XF Analyzer in mouse myometrial smooth muscle cells (mMSMCs). Right, metabolic pathways analyzed. Circled numbers indicate drug injection order. (B) Combined Mitochondrial Stress Test traces for CON (n=8, black) and DIO mMSMCs (n=7, pink). Quantification of (C) area under the curve (AUC), (D) basal respiration rate, (E) ATP-linked respiration, (F) maximal respiration, (G) spare respiratory capacity, (H) non-mitochondrial respiration, (I) proton leak, and (J) coupling efficiency. Results are presented as mean ± SEM. \**P* < 0.05, \*\**P*< 0.01 by Student *t* test. AA, antimycin A; FCCP, trifluoromethoxy carbonyl cyanide phenylhydrazone; OCR, oxygen consumption rate; OG, oligomycin; ROT, rotenone.

### Myometrial cells from obese mice do not increase glycolysis to compensate for reduced mitochondrial respiration

Next, we asked whether mMSMCs from DIO mice can rely on glycolysis for energy production when glucose oxidative phosphorylation is compromised. To evaluate the glycolytic rate, we used a Glycolysis Stress Test assay to determine the ECAR resulting from lactate production (Figure 3A). mMSMCs from DIO uteri exhibited a lower ECAR curve than mMSMCs from CON mice (Figure 3B–C). Moreover, mMSMCs from DIO mice had significantly lower basal glycolysis, glycolytic capacity, and non-glycolytic acidification (Figure 3D–F) than mMSMCs from CON mice. Glycolytic reserve was similar between the groups (Figure 3G). Taken together, these data indicate that, rather than increasing glycolysis to make up for the reduced energy produced during oxidative phosphorylation, mMSMCs from DIO mice have lower glycolysis than MSMCs from CON mice.

**Figure 3:**
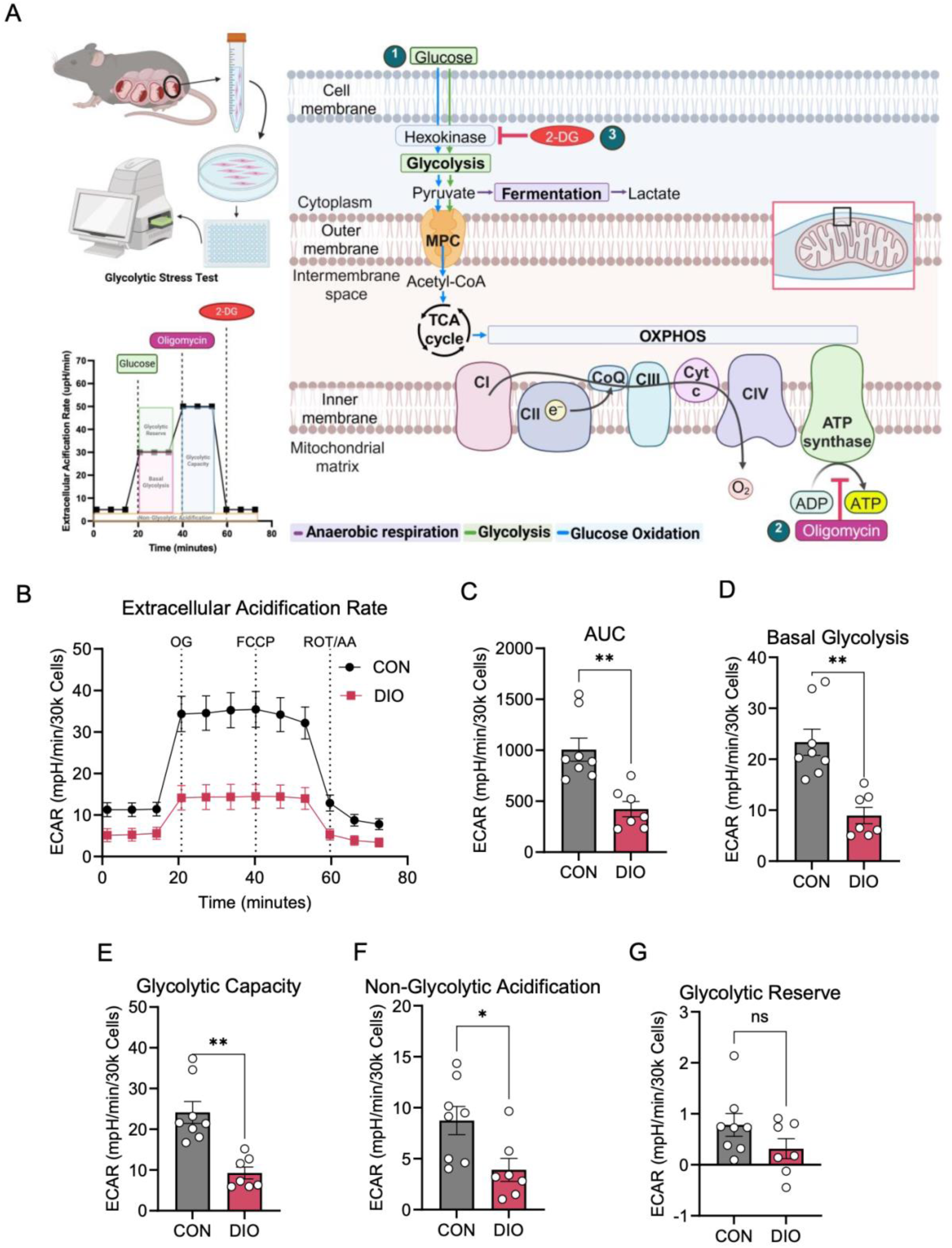
Glycolysis is reduced in DIO myometrial cells. (A) Left, experimental schematic for glycolytic rate with Agilent Seahorse XF Analyzer in mouse myometrial smooth muscle cells (mMSMCs). Right, metabolic pathways analyzed. Circled numbers indicate drug injection order. (B) Combined Glycolytic Stress Test traces for CON (n=8, grey) and DIO mMSMCs (n=7, pink). Quantification of (C) area under the curve (AUC), (D) basal glycolysis, (E) glycolytic capacity, (F) non-glycolytic acidification, and (G) glycolytic reserve. Data are presented as mean ± SEM. \**P* < 0.05, \*\**P*< 0.01, by Student *t* test. ECAR, extracellular acidification rate; GLUC, glucose; OG, oligomycin; 2-DG, 2-deoxy-glucose; ns, not significant.

### Pyruvate transport into mitochondria is disrupted in myometrium from obese mice

Given that mMSMCs from DIO mice had reduced pyruvate utilization and glucose oxidation, we next investigated whether uteri from these mice had altered abundance of glycolytic enzymes and mitochondrial pyruvate transporters (Figure 4A). Abundance of facilitative glucose transporter 1 (GLUT1) and facilitative glucose transporter 3 (GLUT3) protein did not differ between uteri from CON and DIO mice (Figure 4B–C and Supplementary Figure S2A–B), while facilitative glucose transporter 4 (GLUT4) was significantly lower in uteri from DIO compared to CON uteri (Supplementary Figure S2C–D). Additionally, we observed similar abundances of hexokinase-1 (HK1), pyruvate kinase M2 (PKM2), and lactate dehydrogenase A (LDHA) (Figure 4D–G). However, abundances of the mitochondrial pyruvate carrier subunits MPC1 and MPC2 were significantly lower in DIO uteri (Figure 4D, H, I). Although abundance of phosphorylated (inactive) pyruvate dehydrogenase (PDH) relative to total PDH was similar in uteri from CON and DIO mice (Supplementary Figure S2E–G), total PDH was significantly lower in DIO uteri (Figure 4D, J). Collectively, these data suggest that reduced pyruvate transport and its subsequent conversion to acetyl-CoA by PDH contribute to reduced pyruvate flux into the tricarboxylic acid cycle in uteri from DIO mice

**Figure 4:**
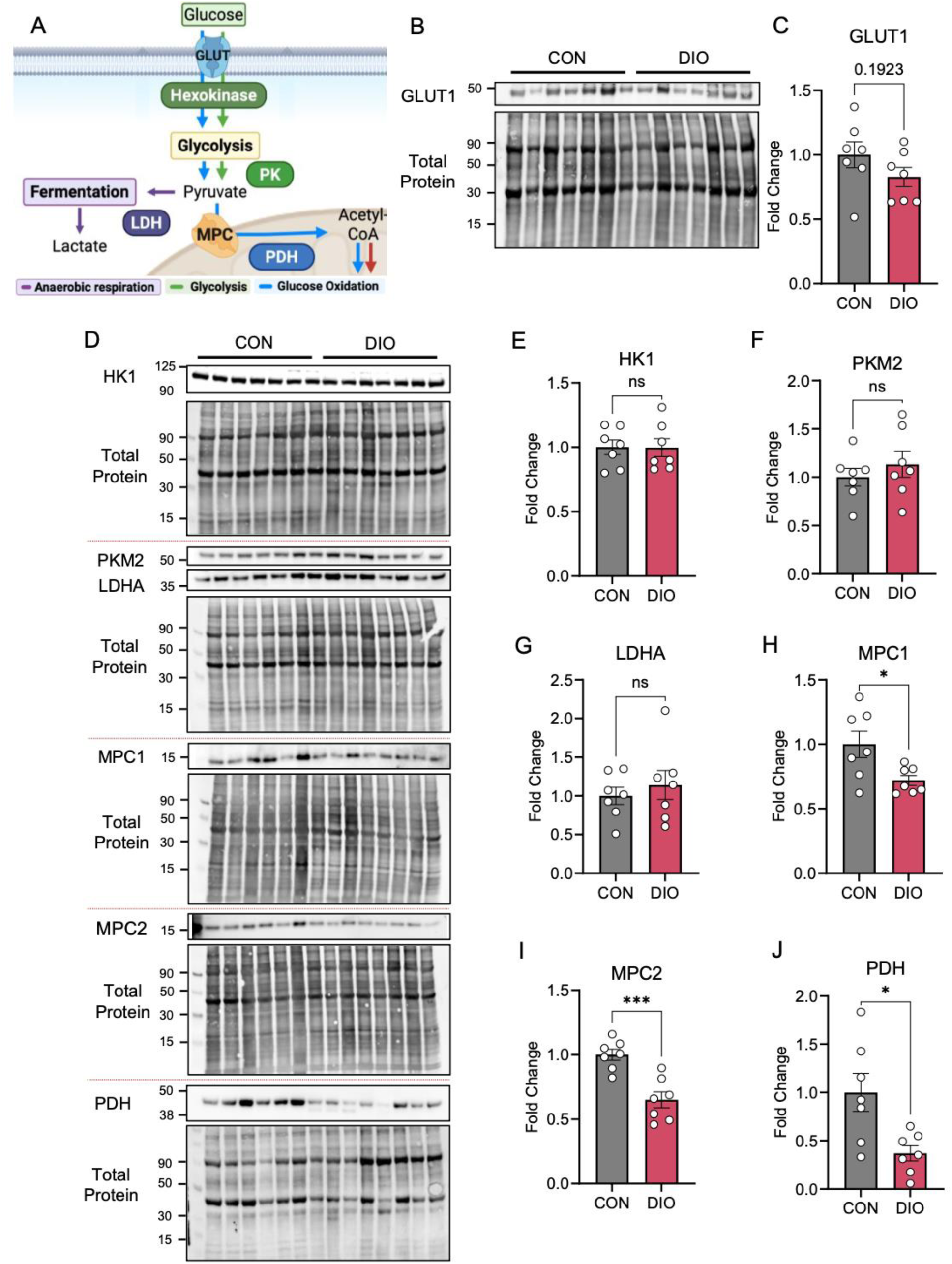
Mitochondrial pyruvate carrier and pyruvate dehydrogenase abundance are decreased in DIO myometrium. (A) Schematic of glucose conversion to pyruvate and acetyl-CoA. (B, D) Representative Western blots probed with antibodies to facilitative glucose transporter 1 (GLUT1), hexokinase 1 (HK1), pyruvate kinase M2 (PKM2), lactate dehydrogenase A (LDHA), mitochondrial pyruvate carrier subunit 1 (MPC1), mitochondrial pyruvate carrier subunit 2 (MPC2), and pyruvate dehydrogenase (PDH), with corresponding total protein blots. Quantification of (C) GLUT1, (E) HK1, (F) PKM2, (G) LDHA, (H) MPC1, (I) MPC2, and (J) total PDH relative to total protein. Data are presented as mean ± SEM. \**P* < 0.05, \*\*\**P* < 0.001 by Student *t* test. ns, not significant.

### Myometrium from obese mice has compromised mitochondria

Changes in mitochondrial function and energy utilization can be linked to shifts in mitochondrial homeostasis. To determine whether mitochondrial homeostasis was affected in DIO uteri, we first assessed mitochondrial morphology and ultrastructure by transmission electron microscopy (Figure 5A). Qualitatively, mitochondria from CON uteri had dense, regular, and tightly packed cristae (Figure 5B). In contrast, mitochondria from DIO uteri had irregular cristae and disrupted matrices. In quantitative analysis, both matrix density and cristae score were significantly lower in DIO mitochondria than in CON mitochondria (Figure 5C–D). Although mitochondrial length and perimeter were similar between groups (Figure 5E–F), DIO mitochondria had significantly greater width, area, circularity, and roundness (Figure 5G–J). Furthermore, mitochondria from DIO uteri aggregated farther from the plasma membrane, had greater mitochondria-endoplasmic reticulum (ER) contact length, and were frequently in proximity to swollen ER (Figure 5K–M). Collectively, these structural and spatial differences suggest that mitochondrial homeostasis is dysregulated in the DIO myometrium.

**Figure 5:**
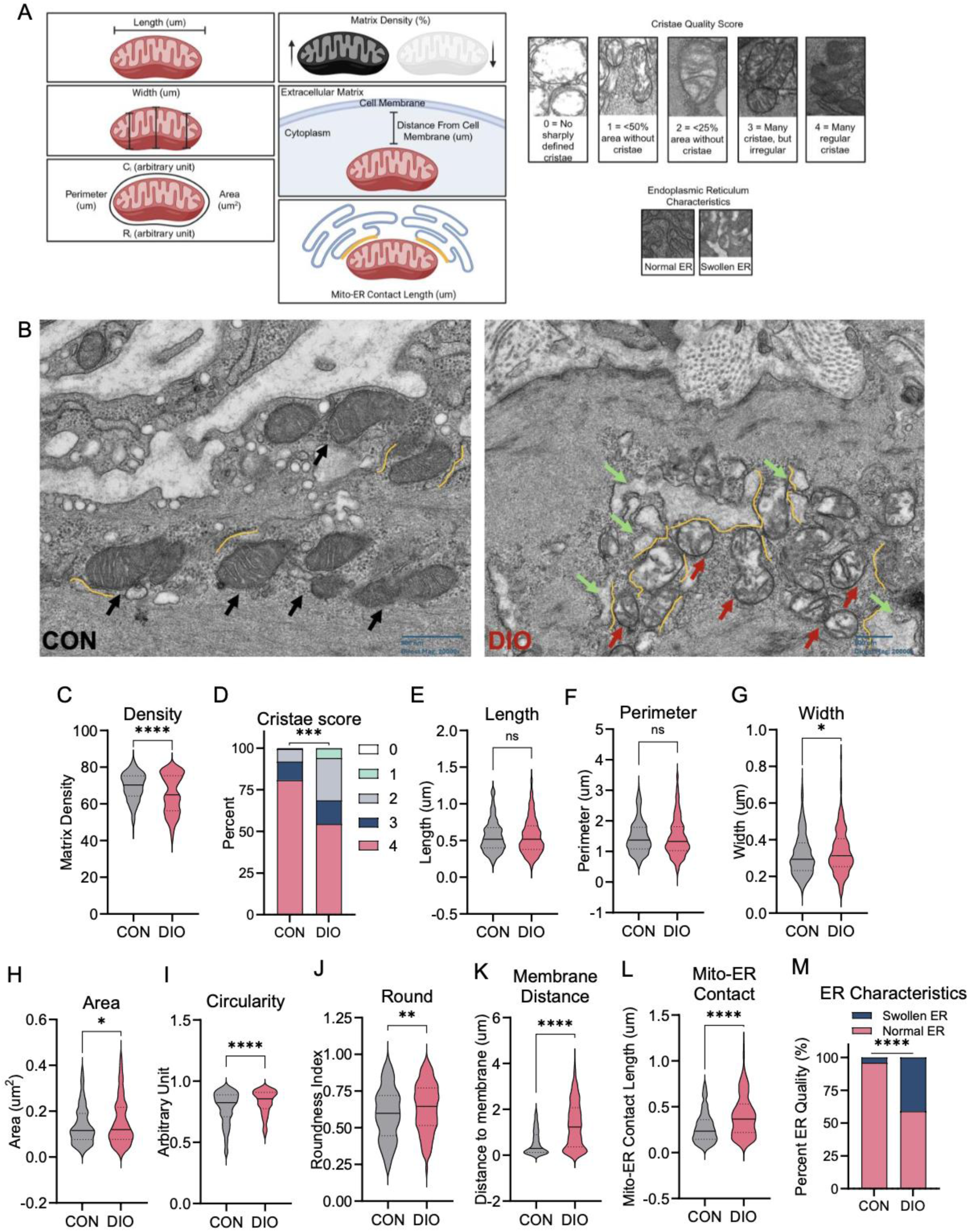
Mitochondria from DIO myometrial tissue have disrupted morphology and ultrastructure. (A) Schematic of transmission electron microscopy (TEM) image analysis. (B) Representative TEM images (scale bar: 500 nm; direct magnification: 20000x) of CON (N=4 mice, black) and DIO (N=4 mice, red) myometrium. CON mitochondria (black arrows), DIO mitochondria (red arrows), swollen endoplasmic reticulum (ER, green arrows), and mitochondria-ER contact length (orange lines) are shown. Quantification of (C) mitochondrial matrix density, (D) cristae score, (E) length, (F) perimeter, (G) width, (H) area, (I) circularity index, (J) roundness index, (K) distance from plasma membrane, (L) mitochondria-ER contact length, and (M) ER characteristics. Data are presented as mean ± SEM. \**P* < 0.05, \*\**P*< 0.01, \*\*\*\**P*< 0.0001 by Student *t* test. ns, not significant.

### Uteri from obese mice have increased mitophagy and decreased mitochondrial fission

Multiple processes maintain mitochondrial homeostasis. Biogenesis generates new mitochondria, mitophagy removes defective mitochondria, and fusion and fission are important for mitochondrial quality control. Additionally, mitochondria are often reversibly tethered to the ER at regions known as mitochondrial-associated membranes (MAMs), which are initiation sites for mitochondrial fusion, fission, and mitophagy. Given our transmission electron microscopy data showing increased mitochondria-ER contact length and ER swelling (Figure 5L–M), we assessed whether MAM protein expression was altered in DIO uteri (Figure 6A). Western blot analysis (Figure 6B) revealed that the mitophagy-associated proteins BCL-2 interacting protein 3 (BNIP3) and microtubule-associated proteins 1A/1B light chain 3B (LC3B-I and LC3B-II), which are involved in degradation of damaged mitochondria, were significantly elevated in DIO uteri (Figure 6C–E). Expression of PTEN-induced kinase 1 (PINK1), which identifies damaged mitochondria and initiates the canonical Parkin-mediated mitophagy pathway, was similar between groups (Figure 6F). Conversely, FUN14 domain containing 1 (FUNDC1), a MAM protein that promotes mitochondrial fission and Parkin-independent mitophagy in response to mitochondrial stress, was significantly decreased in DIO uteri (Figure 6G). Therefore, although both BNIP3 and FUNDC1 can mediate receptor-dependent mitophagy, they changed in opposite directions in DIO uteri. Taken together, these data suggest that DIO is associated with increased BNIP3-associated mitophagy signaling in the uterus, but reduced FUNDC1-dependent mitochondria-ER signaling which, in turn, may impair fission at MAMs. Additionally, abundance of the mitochondrial import receptor translocase of outer mitochondrial membrane 20 (TOM20), which is commonly used as an index of mitochondrial network mass, was significantly lower in DIO uteri than in CON uteri (Figure 6B and 6H), suggesting a possible reduction in mitochondrial capacity.

**Figure 6:**
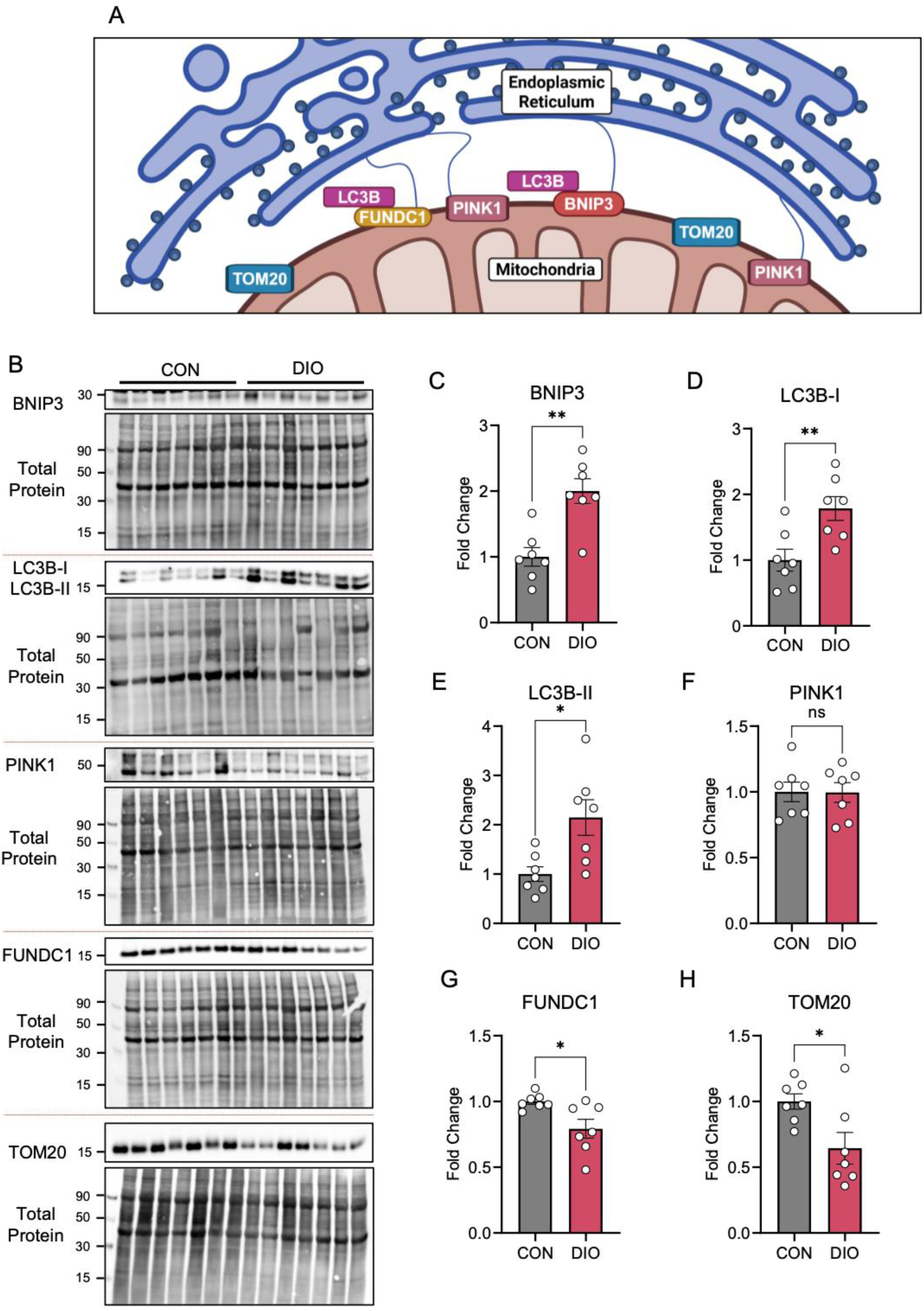
Mitophagy is increased and mitochondria-ER signaling is decreased in DIO myometrium. (A) Schematic of mitophagy and mitochondria-ER associated proteins. (B) Representative Western blots probed with antibodies to BCL2 interacting protein 3 (BNIP3), microtubule-associated protein 1 light chain 3B (LC3B-I and LC3B-II), PTEN-induced kinase 1 (PINK1), FUN14 domain containing 1 (FUNDC1), and translocase of outer mitochondrial membrane 20 (TOM20), with corresponding total protein blots. The total protein blot for FUNDC1 is the same blot for PKM2 and LDHA in Figure 4. The total protein blot for BNIP3 is the same blot for HK1 in Figure 4. Quantification of (C) BNIP3, (D) LC3B-I, (E) LC3B-II, (F) PINK1, (G) FUNDC1, and (H) TOM20 relative to total protein. Data are presented as mean ± SEM. \**P* < 0.05, \*\**P*< 0.01 by Student *t* test. ns, not significant.

Because FUNDC1 drives mitochondrial fission by preferentially recruiting dynamin-related protein 1 (DRP1) in settings of mitochondrial stress, we asked whether mitochondrial dynamics was disrupted in uteri from DIO mice. Normally, fusion, which occurs when two adjacent mitochondria join, and fission, which separates one mitochondrion into two (Figure 7A), counterbalance each other. Inactivation of one leads to unopposed action by the other, resulting in structural and functional imbalance. Mitofusins (MFN1/2) and optic atrophy 1 (OPA1) proteins mediate fusion at the outer and inner mitochondrial membranes, respectively. Conversely, DRP1 is a master regulator of mitochondrial fission. Mitochondrial fission factor (MFF) resides within the outer mitochondrial membrane and is a primary receptor for DRP1, and mitochondrial fission 1 protein (FIS1) is indirectly involved in fission via DRP1 binding. In Western blot analyses (Figure 7B), abundances of the mitochondrial fusion proteins MFN1, MFN2, and OPA1 were similar in DIO and CON uteri (Figure 7C–E). In contrast, the mitochondrial fission proteins DRP1 and MFF were significantly decreased in DIO uteri (Figure 7F and G), and FIS1 abundance was similar between groups (Figure 7H). Taken together, these data suggest a significant decrease in mitochondrial fission machinery in DIO uteri, which, without a compensatory decrease in fusion, shifts the mitochondrial dynamics toward a fusion-dominant state.

**Figure 7:**
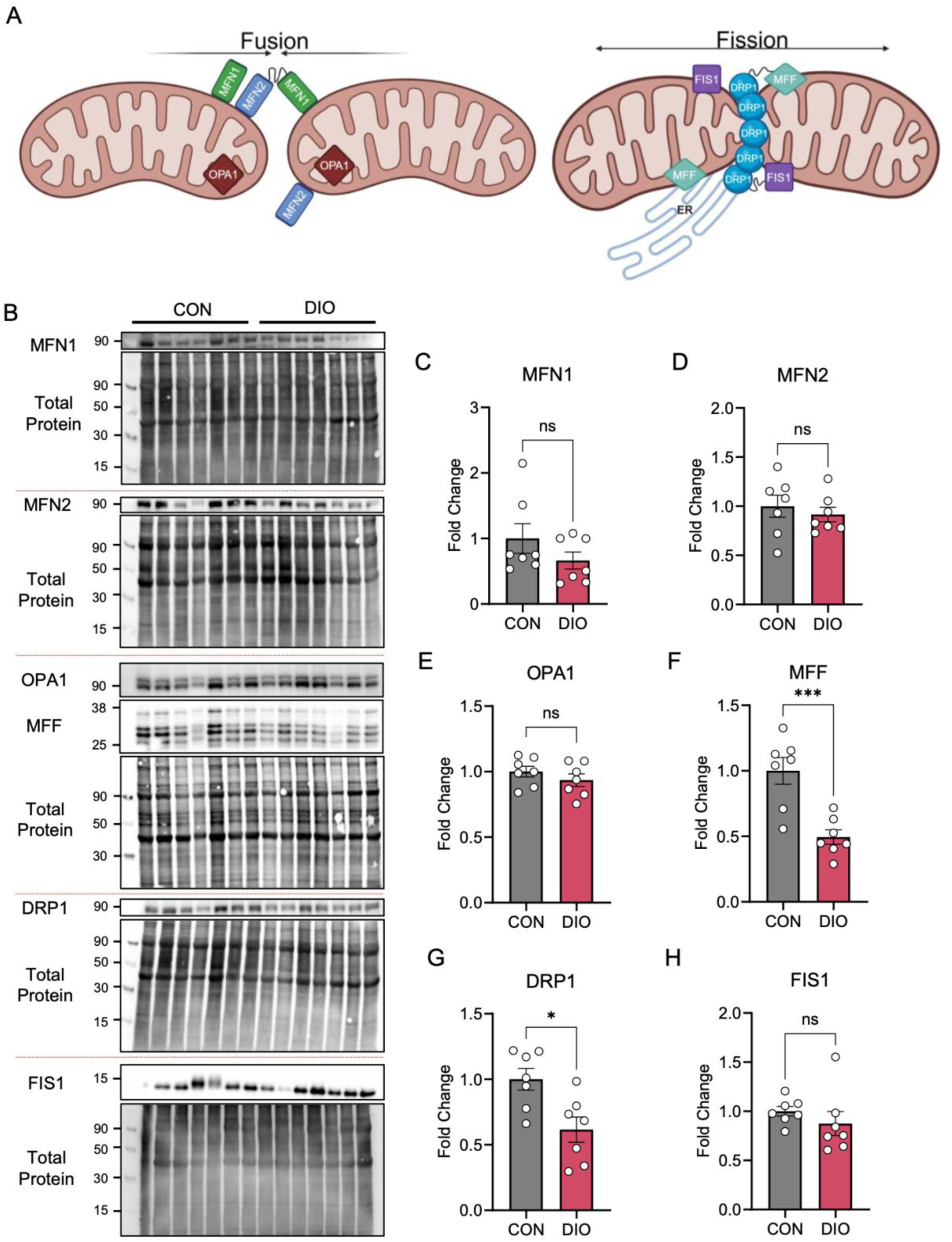
Mitochondrial fission is decreased in DIO myometrium. (A) Schematic of fusion and fission with respective proteins. (B) Representative Western blots probed with antibodies to mitofusin 1 (MFN1), mitofusin 2 (MFN2), optic atrophy 1 (OPA1), mitochondrial fission factor (MFF), dynamin-related protein 1 (DRP1), and mitochondrial fission 1 (FIS1), with corresponding total protein blots. Quantification of (C) MFN1, (D) MFN2, (E) OPA1, (F) MFF, (G) DRP1, and (H) FIS1 relative to total protein. Data are presented as mean ± SEM. \**P* < 0.05, \*\*\**P*< 0.001 by Student *t* test. ns, not significant.

### Exposure to exogenous fatty acids decreases mitochondrial fission proteins in human and mouse myometrial cells

Because we previously found an accumulation of medium-chain fatty acids in DIO uteri^21^, and here observed disruption of mitochondrial dynamics (Figure 7), we asked whether increased fatty acids impaired mitochondrial fission. In humans, plasma concentrations of some of the major fatty acids range between 100 µM and 5 mM^26^. We thus treated primary mMSMCs from CON and DIO mice for 18 hours with 250 µM of the medium-chain fatty acid octanoic acid (C8:0) (Figure 8A) and examined abundances of mitochondrial fission and fusion proteins by Western blot (Figure 8B). Abundance of OPA1, MFF, DRP1 and FIS1 was decreased in mMSMC from DIO compared to CON (Figure 8C-F). Treatment with octanoic acid did not affect abundance of OPA1 (Figure 8C), MFF (Figure 8E), and FIS1 (Figure 8F). However, octanoic acid treatment caused a significant decrease in abundance of DRP1 in CON mMSMCs, approaching levels observed in vehicle-treated DIO cells (Figure 8E). Treatment with octanoic acid did not significantly alter OPA1, MFF, DRP1 and FIS1 in DIO mMSMC compared to vehicle (Figure 8C-F). Collectively, these results indicate that exposure to exogenous octanoic acid, a medium-chain fatty acid, directly downregulate DRP1, a key regulator of fission, leading to impaired mitochondrial fission in murine myometrial cells.

**Figure 8:**
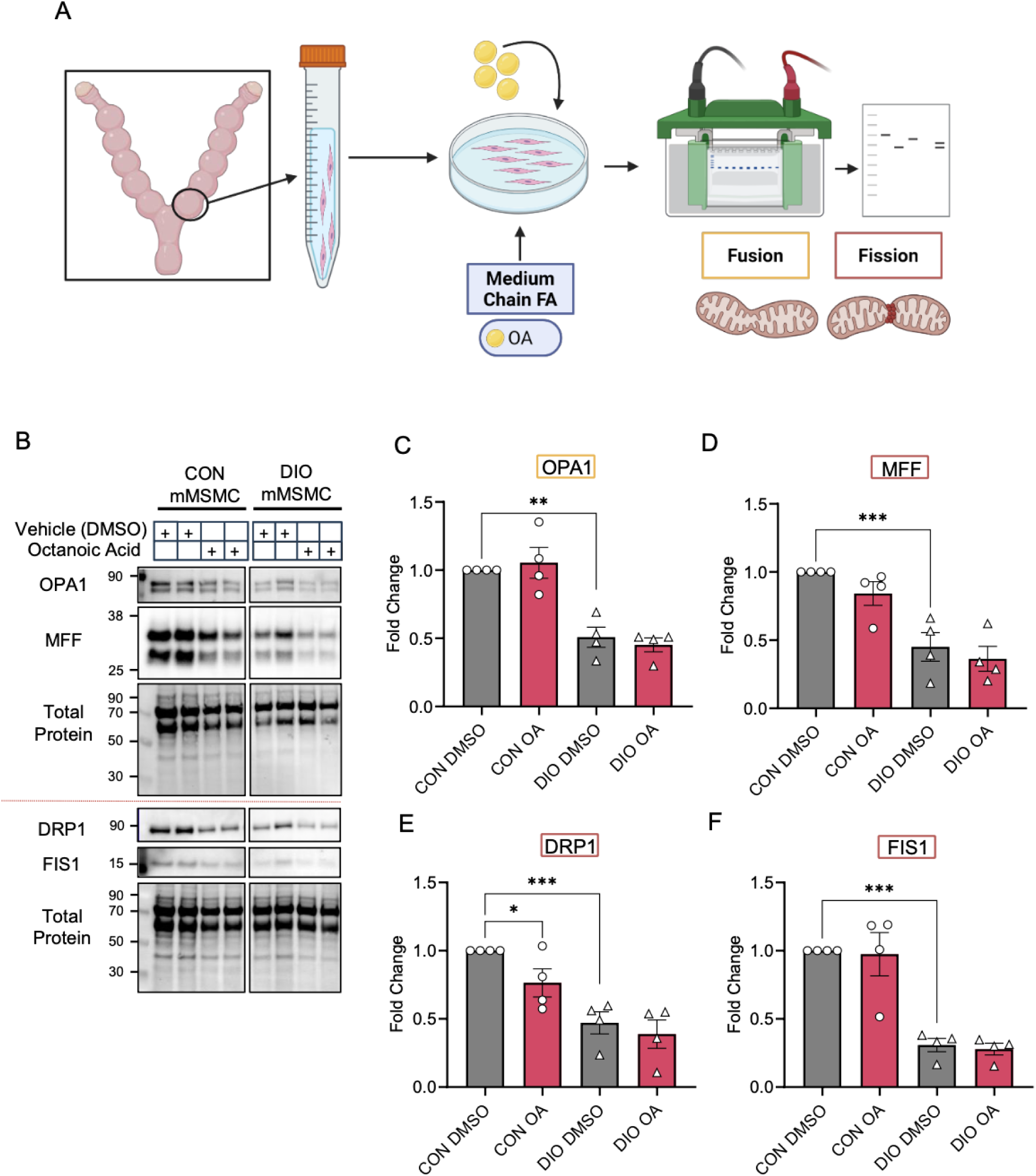
Exposure to medium-chain fatty acid octanoic acid decreases mitochondrial fission in CON myometrial cells. (A) Experimental schematic of *in vitro* treatment of CON murine primary myometrial smooth muscle cells (CON mMSMCs) with vehicle (DMSO) (n=4, grey in C–F) or 250 µM octanoic acid (OA) (n=4, pink in C–F) and of DIO mMSMCs with vehicle (DMSO) (n=4, grey in C–F) or 250 µM octanoic acid (OA) (n=4, pink in C–F). (B) Representative Western blots probed with antibodies to optic atrophy 1 (OPA1), dynamin-related protein 1 (DRP1), mitochondrial fission factor (MFF), and mitochondrial fission 1 (FIS1), with corresponding total protein blots. Quantification of (C) OPA1, (D) MFF, (E) DRP1, and (F) FIS1 relative to total protein. Data are presented as mean ± SEM. \**P* < 0.05 by one-way ANOVA with Holm-Sidak multiple comparison test. ns, not significant.

To establish the clinical and biological relevance of our murine data to humans, we next assessed whether a comparable relationship between medium-chain fatty acids and mitochondrial dynamics exists in human myometrial cells. Similar to the murine experiments, primary human myometrial cells (hMSMCs) isolated from biopsies collected at the time of term, scheduled (unlabored) cesarean delivery were incubated for 18 hours with 250 µM of octanoic acid, followed by western blot analysis (Figure 9A–B). Treatment of hMSMCs octanoic acid did not significantly alter protein abundance of the fusion protein OPA1 (Figure 9C). Octanoic acid treatment significantly decreased the abundance of the fission proteins MFF and DRP1, though FIS1 remained unchanged (Figure 9D–F). These data confirm that medium-chain FA exposure disrupts mitochondrial fission machinery in human myometrial cells.

**Figure 9:**
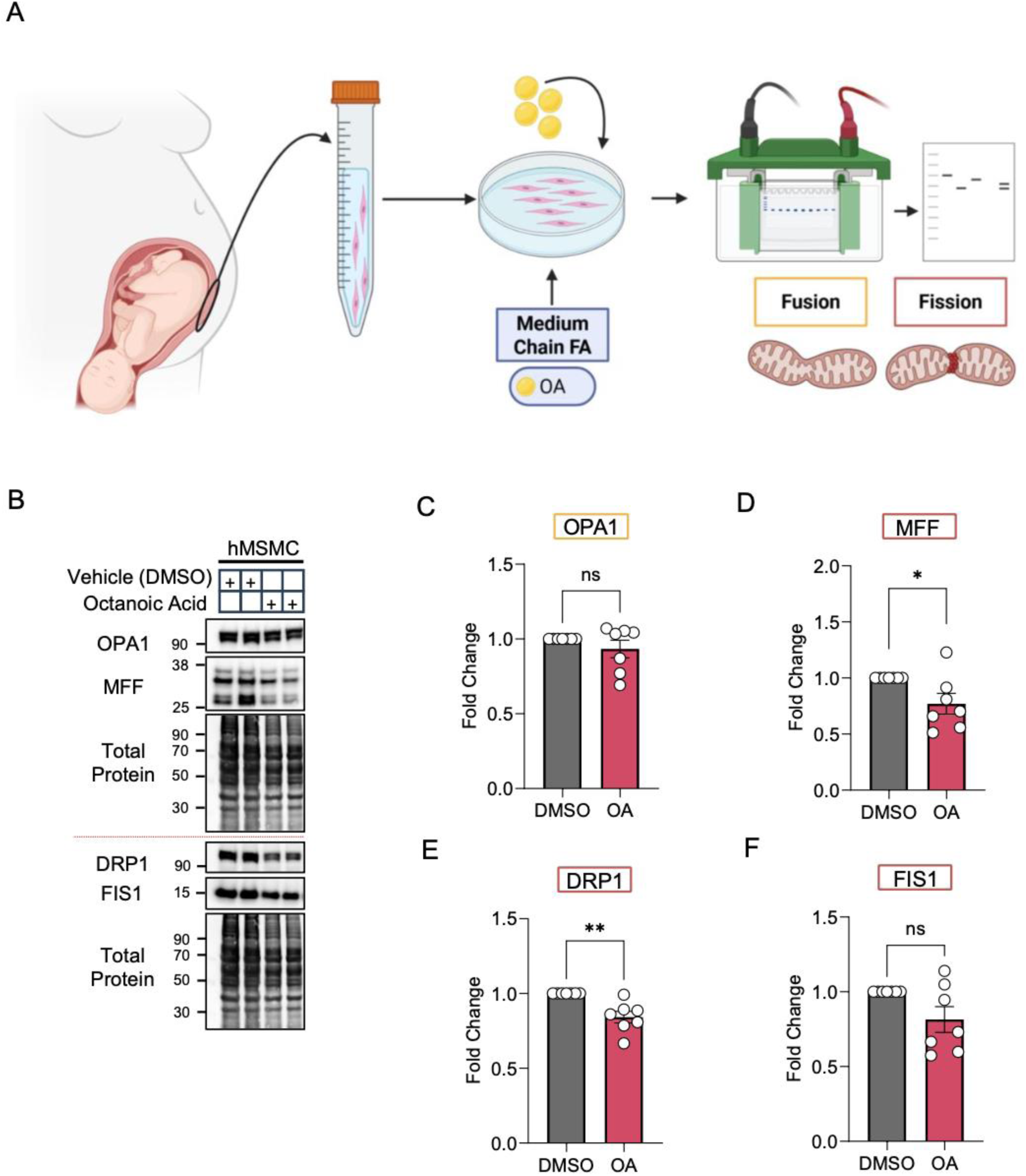
Exposure medium-chain fatty acid octanoic acid decreases mitochondrial fission in human myometrial cells. (A) Experimental schematic of *in vitro* treatment of human primary myometrial smooth muscle cells with vehicle (DMSO) (n=4, grey in C–F) or 250 µM octanoic acid (OA) (n=4, pink in C–F). (B) Representative Western blots probed with antibodies to optic atrophy 1 (OPA1), mitochondrial fission factor (MFF), dynamin-related protein 1 (DRP1), and mitochondrial fission 1 (FIS1), with corresponding total protein blots. Quantification of (C) OPA1, (D) MFF, (E) DRP1, and (F) FIS1 relative to total protein. Data are presented as mean ± SEM. \**P* < 0.05 by one-way ANOVA with Holm-Sidak multiple comparison test. ns, not significant.

### Myometrial cells from patients with obesity have lower mitochondrial respiration rate than cells from patients without obesity

Finally, we asked whether BMI was associated with decreased mitochondrial energy production in the human myometrium. To test this, we obtained hMSMCs from patients with BMI values between 20 and 74 kg/m^2^ (Figure 10A and Table 1) and analyzed them with the Seahorse XF Cell Mito Stress Test, as described^15^. The OCR curve was lower in hMSMCs from patients with BMI ≥ 30 kg/m^2^ than in hMSMCs from patients with BMI < 30 kg/m^2^ (Figure 10B), though the areas under the curve were not significantly different between the two groups (Figure 10C). Analysis of ECAR after addition of oligomycin and OCR/ECAR ratios at baseline, after oligomycin, and at maximal respiration did not significantly differ between groups (Supplementary Figure S4). Given the wide range of BMI values in the cohort, we performed a linear regression to assess the relationship between BMI and individual mitochondrial function parameters. Basal respiration, ATP linked respiration, non-mitochondrial respiration, and proton leak did not significantly differ by BMI. However, spare capacity significantly decreased with increasing BMI, showing a 0.38 pmol/min decrease in OCR with every point increase in BMI (Figure 10D). We performed adjusted analyses including age in the model, as greater age has a deleterious effect on muscle mitochondrial function^27^ and differed between our obese and non-obese groups, but this did not change the relationship and was not kept in the final model. Maximal respiration decreased, though not significantly with BMI (Figure 10E). Taken together, these data suggest that, consistent with our mouse data, maternal obesity in humans is associated with a reduced capacity of primary hMSMCs to increase oxygen consumption beyond basal levels.

**Figure 10:**
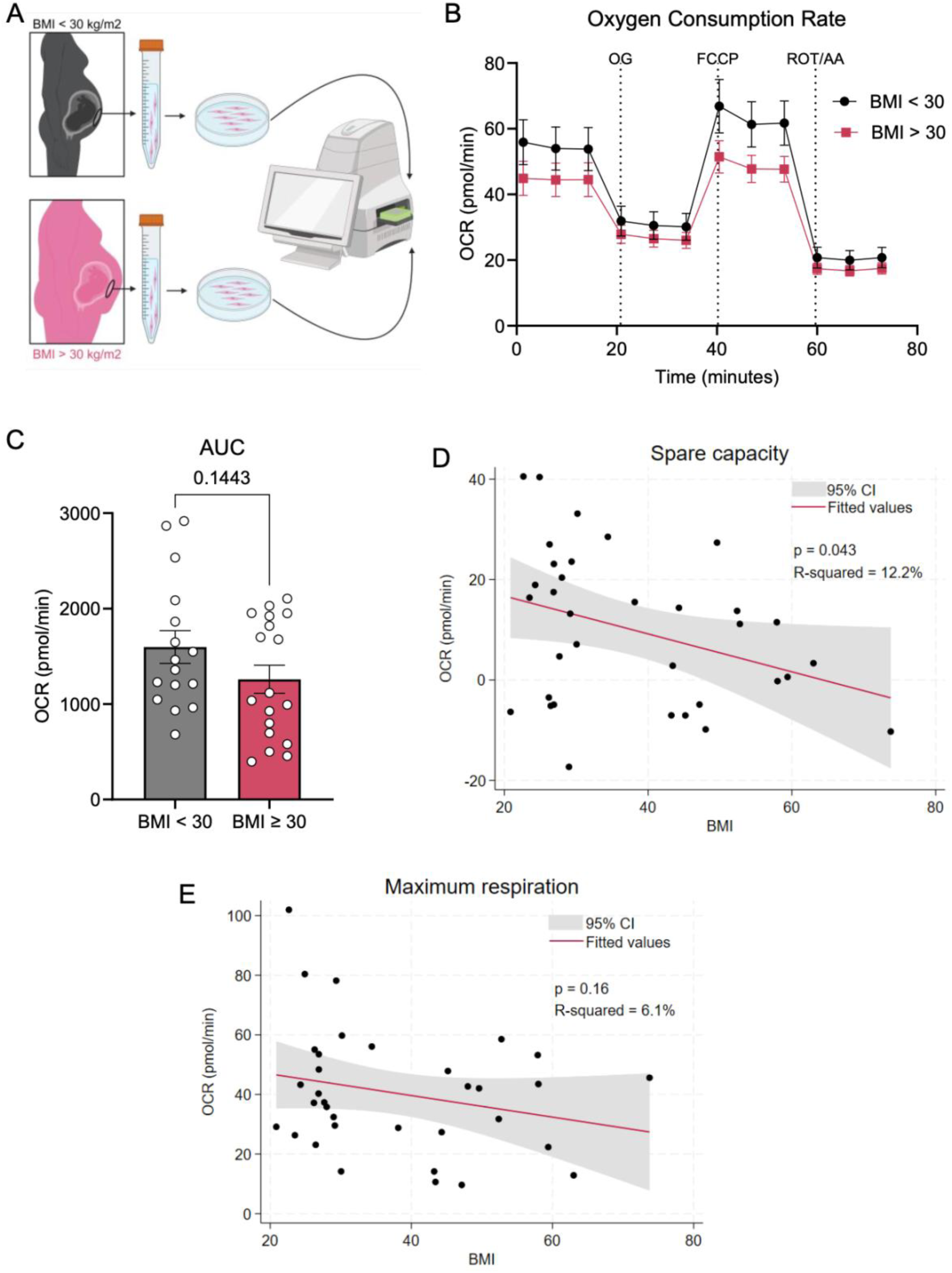
Human myometrial smooth muscle cells from patients with higher BMI have reduced oxygen consumption rate and mitochondrial maximal respiration. A) Experimental schematic for mitochondrial respiration with Agilent Seahorse XF Analyzer in human primary myometrial smooth muscle cells (hMSMCs) from patients with body mass index (BMI) < 30 kg/m^2^ or BMI ≥ 30 kg/m^2^. (B) Combined Mitochondrial Stress Test traces for hMSMCs from patients with BMI < 30 kg/m^2^ (n=16, black) or BMI ≥ 30 kg/m2 (n=18, pink). (C) Quantification of area under the curve (AUC). Linear regression was used to assess the relationship between maternal BMI and (D) spare respiratory capacity or (E) maximal respiration. Fitted lines (red) with 95% confidence intervals (gray) for the predicted means are presented. Data are presented as mean ± SEM. Statistical analysis by one-way ANOVA with Holm-Sidak multiple comparison test. AA, antimycin A; ECAR, extracellular acidification rate; FCCP, trifluoromethoxy carbonyl cyanide phenylhydrazone; OCR, oxygen consumption rate; OG, oligomycin; ROT, rotenone.

## DISCUSSION

Here, we show that maternal obesity disrupts mitochondrial fission and inhibits glucose oxidation in the uterus at term. We previously showed that obesity inhibits spontaneous uterine contractility, upregulates fatty acid oxidation, and results in accumulation of medium-chain acylcarnitines in the murine uterus^21^. We propose that this accumulation of fatty acids disrupts mitochondrial dynamics, resulting in a decrease in fission-associated proteins and an increase in mitophagy. This leads to a decrease in mitochondrial pyruvate carriers on the mitochondrial membrane, resulting in decreased pyruvate utilization, dysregulation of glucose oxidation, and reduced energy generation in the myometrium. These effects may impair myometrial contractility and contribute to poor labor outcomes.

Data regarding mitochondrial function in the myometrium are limited. Gam et al. compared myometrial tissue from elective cesarean deliveries of women with and without obesity and reported no significant mitochondrial dysfunction^28^. This is in contrast our finding that uteri from DIO mice had compromised mitochondrial structure and function. However, Gam et al. only assessed oxidative capacity in isolated mitochondria and did not assess mitochondrial dynamics. Nonetheless, their paper shows a decrease in respiratory control ratio in myometrial tissue from women with obesity, suggesting decreased efficiency of ATP generation in the isolated mitochondria^28^. Furthermore, although Gam et al. did not report a difference in mitochondrial morphology via electron microscopy, the images were only qualitatively analyzed. Instead, our rigorous, blinded, quantitative analysis of transmission electron microscopy images revealed that mitochondria in DIO mouse uteri had greater variability in shape and size, less dense matrices, and fewer cristae than those in CON uteri. Furthermore, the abundance of the outer mitochondrial membrane protein TOM20 was significantly lower in uterine tissue from DIO mice than in tissue from CON mice. This finding indicates a disrupted mitochondrial network and suggests that mitochondrial homeostasis and dynamics are impaired in DIO uteri.

Fusion and fission occur continuously to maintain mitochondrial homeostasis, and disruption of either process impairs cellular energy production. For example, DRP1 inhibition resulted in decreased mitochondrial respiration, impaired function, and decreased mitochondrial DNA copy number in HeLa cells^29,30^. In skeletal muscle, DRP1-deficient mitochondria increase in size and are energetically dysfunctional, resulting in increased mitochondrial calcium uptake and muscle atrophy^31^. Studies in skeletal muscle have shown that excess palmitate can result in mitochondrial fragmentation and increased fission machinery protein abundance, leading to smaller mitochondria in mice with genetic and diet-induced obesity^18^. Additionally, ceramide accumulation results in elevated mitochondrial fission and increased DRP1 expression^32^. In both our DIO model and in fatty acid-treated cells *in vitro*, we observed a decrease in DRP1 and its partners MFF and FIS1. This could be a compensatory mechanism in response to increased fatty acid oxidation to prevent further mitochondrial loss^33^. Such a mechanism can be explored in future studies.

In DIO uteri, we noted a reduction in fission proteins (DRP1, MFF, FIS1) without a concomitant decrease in fusion protein abundance. Although this might be expected to shift mitochondria toward a more fusion-dominated, elongated network^34^, we instead observed more rounded mitochondria, which is consistent with fragmentation and swelling rather than elongation. One possible explanation is that the disrupted ER-contacts and fission signaling at MAMs results in an inability to clear the damaged mitochondria, despite an overall bias in machinery towards fusion^35^. Another possibility is that the fission machinery decreased in response to pre-existing mitochondrial fragmentation or damage rather than caused it. Finally, the total protein abundance may reflect functional changes in fission and fusion as these proteins are also regulated by posttranslational modification and subcellular localization^36^. Nonetheless, the observed mitochondrial morphological changes, and the changes in fission-associated and MAM proteins, together support a change in mitochondrial dynamics associated with DIO and exogenous fatty acid exposure. Future work should examine the direct functional effects of changes in mitochondrial dynamics and longitudinally evaluate mitochondrial dynamics upon myometrial cell exposure to fatty acids.

Disruption of the fusion-fission balance can also elicit mitophagy, a form of specialized autophagy that removes damaged mitochondria^37^. Mitophagy and ER signaling are likely dysregulated in DIO uteri, as we observed altered mitochondrial localization, increased mitochondria-ER contact length, and ER swelling. Mitophagy can proceed via several pathways, including PINK1/parkin-mediated mitophagy, hypoxia-mediated FUNDC1 activation, and the BNIP3 receptor pathway^38^. Our data implicate the latter two, which both utilize LC3B. During mitophagy, mitochondria are in proximity to the ER, and mitochondria-ER contact sites (MERCs) help maintain mitochondrial homeostasis as mitophagy progresses^38^. MERCs, which also regulate cellular metabolism, calcium signaling, and organelle homeostasis^39^, are mediated by mitochondria-associated ER membrane proteins, which help maintain mitochondrial and ER integrity through processes such as mitophagy^40,41^. Disruption of MERC signaling is linked to skeletal muscle dysfunction in obesity^42^. In the myometrium of obese mice, it is thus likely that MERC signaling is decreased.

Data are limited regarding myometrial energy metabolism during parturition, particularly in the setting of maternal obesity. In skeletal and cardiac muscle, obesity upregulates fatty acid β-oxidation, inhibits glucose oxidation, and leads to dysfunctional contractility due to failure to metabolically switch between substrates^43–45^. Limited previous research suggests that, in women without obesity, the myometrium uses glucose and glycogen as its main sources of energy in preparation for labor^15,46–49^. In primary hMSMCs isolated from women without obesity at term, the addition of UK5099, which inhibits glucose oxidation by blocking the mitochondrial pyruvate carrier, reduced basal OCR and spare respiratory capacity^15^. In contrast, etomoxir, which inhibits fatty acid oxidation by blocking carnitine palmitoyltransferase, had no effect on OCR. This finding suggests that human myometrial cells use glucose as their primary source of energy and are not acutely metabolically flexible. Here, we show that obesity decreases the ability of the myometrium to utilize carbohydrates, its preferred energy source, for oxidative phosphorylation. This likely leads to the observed decrease in mitochondrial energy generation observed in obese mice. We also show that this myometrial phenotype is conserved, as increasing BMI in humans was associated with decreased capacity for myometrial cells to increase oxygen consumption beyond basal levels.

The mitochondrial phenotype we observed in permeabilized myometrial fibers differed from that observed in isolated mMSMCs. In fibers, maternal obesity led to impaired pyruvate-driven oxidative phosphorylation but no effects on fatty acid oxidation and RCR. This suggests an isolated defect in carbohydrate handling rather than global impairment of mitochondrial coupling or structure. In contrast, isolated primary mMSMCs from DIO mice exhibited reductions in basal, ATP-linked, and maximal respiration. This discrepancy likely reflects a key difference between the two assays. Whereas the permeabilized fiber protocol isolates mitochondrial flux through a single, defined substrate pathway, the Mito Stress Test captures an integrated readout of respiration utilizing all substrates available to the cell, including exogenous components of the assay media and endogenous lipid stores accumulated as a result of DIO. Therefore, dysfunction in a single substrate pathway would manifest as a broader OCR reduction in isolated cells. Future studies using substrate-specific inhibitors (e.g., etomoxir, UK5099) in the intact-cell Seahorse assay would help determine the relative contribution of endogenous lipid versus carbohydrate oxidation to the broader bioenergetic defect observed in DIO mMSMCs. Together, our data suggest that the mitochondrial dysfunction observed in DIO myometrium primarily affects carbohydrate oxidative phosphorylation. This conclusion is consistent with our prior work showing that myometrial bioenergetics and contractility primarily depend on glucose oxidation and rely minimally on fatty acid oxidation under normal physiological conditions^15^.

DIO myometrial cells did not have a concomitant increase in glycolysis despite increased abundances of GLUT1 and GLUT4, suggesting more glucose entering the cytosol. The decrease in ECAR in isolated DIO myometrial cells coupled with the unchanged abundances of HK1, PKM2, and LDHA in DIO uterine tissue could be due to differences in cell versus tissue capacity for glycolysis. These differences could also be due to glucose and fatty acids competing for oxidation in the Randle cycle^50,51^. During the Randle cycle, elevated fatty acid oxidation rapidly increases mitochondrial acetyl-CoA/CoA and NADH/NAD^+^ ratios, resulting in inhibition of PDH and subsequent accumulation of citrate and increased ATP, which can inhibit glycolytic enzymes^51,52^. This is consistent with our observed decrease in total PDH protein abundance here as well as the reported increase in citrate and ATP/ADP ratio in DIO uteri^21,53^. These changes could result in suppression of glycolysis in DIO myometrial cells and rerouting of glucose to other metabolic pathways, such as glycogen storage, the pentose phosphate pathway, or the hexosamine biosynthesis pathway^51,54^. In the future, metabolite tracing experiments could be performed to determine the fate of glucose in myometrial cells in the setting of maternal obesity.

Previous research suggests that an imbalance in nutrient supply and demand can affect mitochondrial function in muscle^55^. In skeletal muscle, excess fatty acids from prolonged exposure to diets high in fat can cause incomplete β-oxidation, which leads to accumulation of fatty acid intermediates that disrupt both glucose oxidation and insulin signaling^45^. Accumulation of fatty acid intermediates can also lead to mitochondrial electron transport chain failure and reduced energy generation^56^. Although research is limited regarding the myometrium, metabolomic data indicate that, compared to CON uterine tissue, DIO uterine tissue contains more beta-oxidation intermediates, specifically medium- and long-chain acylcarnitines^21^. Our current data expand upon these findings to show that fatty acid exposure results in mitochondrial dysfunction, decreased energy production, and abnormal glucose utilization in the myometrium. Although we did not assess insulin signaling in the myometrium, we postulate that it is disrupted in the setting of maternal obesity. Further research is needed to test whether impaired insulin signaling contributes to labor dystocia in women with obesity.

Our study extends previous work in several important ways. First, whereas Gam et al. characterized static respiratory function in isolated mitochondria, we assessed mitochondrial fission, fusion, and mitophagy and in an intact cellular environment. Second, we showed that exposure to a physiologically relevant fatty acid concentration is sufficient to reproduce the fission defect observed in the setting of obesity. Third, we report OCR data in primary myometrial cells isolated from 34 women with a wide range of BMI values. Combined with our fatty acid exposure data, this allows us to move from a population-level association between BMI and mitochondrial function to a mechanistic model.

This study has three key limitations and thus raises important new questions. First, DRP1 regulates both mitochondrial and peroxisomal fission^55^, and we did not assess peroxisomal function in DIO uteri. Further studies are needed to assess the effects of obesity on peroxisomal dynamics and function in the myometrium and determine whether oxidation of very long-chain fatty acids in the peroxisome contributes to energy deficiencies. Second, we focused on medium-chain fatty acids based on our previous observations of an accumulation of medium-chain acylcarnitines in the murine uterus from our DIO model^21^. The lower abundance of DRP1 in DIO mMSMCs compared to octanoic acid treated CON mMSMCs may be due to additive effects of many contributing factors and further studies are needed to assess the effects of different length-chain FAs, different saturations of FAs, as well as high levels of sucrose on mitochondrial function and energy production in the uterus. Third, we did not assess generation of reactive oxygen species, which increase in cases of nutrient excess from high-fat, high-sucrose diets, especially after tissue becomes insulin resistant^57^. In muscle from diabetic animals, mitochondrial electron transport chain activity is decreased, and this may be a regulatory mechanism to protect against insulin resistance^58,59^. Moreover, muscle-specific knockouts of either apoptosis-inducing factor or mitochondrial transcription factor A, both of which result in reduced electron transport chain activity, ameliorate insulin sensitivity^58,59^. Future studies are needed to determine whether, in the myometrium of obese animals, the reduction in energy generation resulting from blunted electron transport chain activity reduces reactive oxygen species-induced insulin resistance.

In conclusion, our data show that a mouse model of maternal DIO results in structural and functional mitochondrial disruption in the myometrium at term. Transmission electron microscopy of uterine tissue suggests damage to the mitochondria in DIO mice, along with dysfunctional mitochondria-ER tethering. A significant decrease in mitochondrial fission-associated proteins was observed in DIO myometrial tissue, accompanied by evidence of increased mitophagy. This decrease in fission was also present *in vitro* in primary CON murine myometrial cells and primary human myometrial cells treated with medium-chain fatty acid octanoic acid. These structural changes were accompanied by decreased pyruvate utilization and energy production in the myometrium, confirmed in both tissue and isolated primary cells. This was mediated by reduced pyruvate translocation and utilization via MPC and PDH, respectively. Increasing BMI in a human cohort was associated with reduced myometrial mitochondrial capacity similar to that observed in DIO murine myometrial cells, supporting the translational relevance of these findings across species.

## AUTHOR CONTRIBUTIONS

AIF and KKP conceptualization; AIF funding acquisition; KKP, JLR, AMG, RMG, TAP, WLB, and AIF methodology; KKP, JLR, AMG, TAP and AIF investigation and experimental work; KKP, WLB, and AIF interpretation of results; KKP and AIF writing—original draft; KKP, JLR, AMG, RMG, TAP, WLB, and AIF writing—review and editing. All authors read and approved the final version.

## ACKNOWLEDGEMENTS

We thank Dr. Deborah Frank for invaluable editorial assistance. We thank the Reproductive Specimen Processing and Banking core (ReProBank) for assistance with human specimen procurement. We thank the patients at Barnes Jewish Hospital for being part of this research study. Seahorse was performed and supported by the Washington University Diabetes Research Center (P30DK020579). This was supported by NIH grant R21 HD110610 (A.I.F.).

## REFERENCES

1. Ogden CL, Carroll MD, Kit BK, Flegal KM. Prevalence of childhood and adult obesity in the United States, 2011-2012. Jama. Feb 26 2014;311(8):806–14. doi:10.1001/jama.2014.732

2. Ellis JA, Brown CM, Barger B, Carlson NS. Influence of Maternal Obesity on Labor Induction: A Systematic Review and Meta-Analysis. J Midwifery Womens Health. Jan 2019;64(1):55–67. doi:10.1111/jmwh.12935

3. Alexander JM, McIntire DD, Leveno KJ. Forty weeks and beyond: pregnancy outcomes by week of gestation. Obstet Gynecol. Aug 2000;96(2):291–4. doi:10.1016/s0029-7844(00)00862-0

4. Conner SN, Verticchio JC, Tuuli MG, Odibo AO, Macones GA, Cahill AG. Maternal obesity and risk of postcesarean wound complications. Am J Perinatol. Apr 2014;31(4):299–304. doi:10.1055/s-0033-1348402

5. Garabedian MJ, Williams CM, Pearce CF, Lain KY, Hansen WF. Extreme morbid obesity and labor outcome in nulliparous women at term. Am J Perinatol. Oct 2011;28(9):729–34. doi:10.1055/s-0031-1280852

6. Norman SM, Tuuli MG, Odibo AO, Caughey AB, Roehl KA, Cahill AG. The effects of obesity on the first stage of labor. Obstet Gynecol. Jul 2012;120(1):130–5. doi:10.1097/AOG.0b013e318259589c

7. Frolova AI, Wang JJ, Conner SN, et al. Spontaneous Labor Onset and Outcomes in Obese Women at Term. Am J Perinatol. Jan 2018;35(1):59–64. doi:10.1055/s-0037-1605574

8. Carlson NS, Hernandez TL, Hurt KJ. Parturition dysfunction in obesity: time to target the pathobiology. Reprod Biol Endocrinol. Dec 18 2015;13:135. doi:10.1186/s12958-015-0129-6

9. Burstein E, Levy A, Mazor M, Wiznitzer A, Sheiner E. Pregnancy outcome among obese women: a prospective study. Am J Perinatol. Oct 2008;25(9):561–6. doi:10.1055/s-0028-1085623

10. Wolfe KB, Rossi RA, Warshak CR. The effect of maternal obesity on the rate of failed induction of labor. Am J Obstet Gynecol. Aug 2011;205(2):128.e1-7. doi:10.1016/j.ajog.2011.03.051

11. Hill M, Reed KL, Cohen WR. Oxytocin utilization for labor induction in obese and lean women. J Perinat Med. Nov 2015;43(6):703–6. doi:10.1515/jpm-2014-0134

12. Roloff K, Peng S, Sanchez-Ramos L, Valenzuela GJ. Cumulative oxytocin dose during induction of labor according to maternal body mass index. Int J Gynaecol Obstet. Oct 2015;131(1):54–8. doi:10.1016/j.ijgo.2015.04.038

13. Fyfe EM, Thompson JM, Anderson NH, Groom KM, McCowan LM. Maternal obesity and postpartum haemorrhage after vaginal and caesarean delivery among nulliparous women at term: a retrospective cohort study. BMC Pregnancy Childbirth. Oct 18 2012;12:112. doi:10.1186/1471-2393-12-112

14. Zhang J, Bricker L, Wray S, Quenby S. Poor uterine contractility in obese women. BJOG. Mar 2007;114(3):343–8. doi:10.1111/j.1471-0528.2006.01233.x

15. Prifti KK, Dave RM, Mitchum KT, et al. Energy substrate preference and metabolic flexibility in quiescent and oxytocin-stimulated human myometrial cells. iScience. Jun 19 2026;29(6):116248. doi:10.1016/j.isci.2026.116248

16. Liu YJ, McIntyre RL, Janssens GE, Houtkooper RH. Mitochondrial fission and fusion: A dynamic role in aging and potential target for age-related disease. Mech Ageing Dev. Mar 2020;186:111212. doi:10.1016/j.mad.2020.111212

17. Wai T, García-Prieto J, Baker MJ, et al. Imbalanced OPA1 processing and mitochondrial fragmentation cause heart failure in mice. Science. Dec 04 2015;350(6265):aad0116. doi:10.1126/science.aad0116

18. Jheng HF, Tsai PJ, Guo SM, et al. Mitochondrial fission contributes to mitochondrial dysfunction and insulin resistance in skeletal muscle. Mol Cell Biol. Jan 2012;32(2):309–19. doi:10.1128/MCB.05603-11

19. Jheng HF, Huang SH, Kuo HM, Hughes MW, Tsai YS. Molecular insight and pharmacological approaches targeting mitochondrial dynamics in skeletal muscle during obesity. Ann N Y Acad Sci. Sep 2015;1350:82–94. doi:10.1111/nyas.12863

20. Zheng P, Ma W, Gu Y, et al. High-fat diet causes mitochondrial damage and downregulation of mitofusin-2 and optic atrophy-1 in multiple organs. J Clin Biochem Nutr. Jul 2023;73(1):61–76. doi:10.3164/jcbn.22-73

21. Prifti KK, McCarthy R, Ma X, Finck BN, England SK, Frolova AI. Obese mice have decreased uterine contractility and altered energy metabolism in the uterus at term gestation. Biol Reprod. Jun 10 2024;doi:10.1093/biolre/ioae086

22. Pietka TA, Brookheart RT. Measurement of Mitochondrial Respiration in Human and Mouse Skeletal Muscle Fibers by High-Resolution Respirometry. J Vis Exp. Oct 4 2024;(212)doi:10.3791/66834

23. Neikirk K, Lopez EG, Marshall AG, et al. Call to action to properly utilize electron microscopy to measure organelles to monitor disease. Eur J Cell Biol. Dec 2023;102(4):151365. doi:10.1016/j.ejcb.2023.151365

24. Lam J, Katti P, Biete M, et al. A Universal Approach to Analyzing Transmission Electron Microscopy with ImageJ. Cells. Aug 24 2021;10(9)doi:10.3390/cells10092177

25. Garcia-Flores V, Romero R, Peyvandipour A, et al. A single-cell atlas of murine reproductive tissues during preterm labor. Cell Rep. Jan 31 2023;42(1):111846. doi:10.1016/j.celrep.2022.111846

26. Abdelmagid SA, Clarke SE, Nielsen DE, et al. Comprehensive profiling of plasma fatty acid concentrations in young healthy Canadian adults. PLoS One. 2015;10(2):e0116195. doi:10.1371/journal.pone.0116195

27. Short KR, Bigelow ML, Kahl J, et al. Decline in skeletal muscle mitochondrial function with aging in humans. Proc Natl Acad Sci U S A. Apr 12 2005;102(15):5618–23. doi:10.1073/pnas.0501559102

28. Gam CMBF, Larsen LH, Mortensen OH, et al. Unchanged mitochondrial phenotype, but accumulation of lipids in the myometrium in obese pregnant women. J Physiol. Dec 01 2017;595(23):7109–7122. doi:10.1113/JP274838

29. Benard G, Bellance N, James D, et al. Mitochondrial bioenergetics and structural network organization. J Cell Sci. Mar 1 2007;120(Pt 5):838–48. doi:10.1242/jcs.03381

30. Parone PA, Da Cruz S, Tondera D, et al. Preventing mitochondrial fission impairs mitochondrial function and leads to loss of mitochondrial DNA. PLoS One. Sep 22 2008;3(9):e3257. doi:10.1371/journal.pone.0003257

31. Favaro G, Romanello V, Varanita T, et al. DRP1-mediated mitochondrial shape controls calcium homeostasis and muscle mass. Nat Commun. Jun 12 2019;10(1):2576. doi:10.1038/s41467-019-10226-9

32. Smith ME, Tippetts TS, Brassfield ES, et al. Mitochondrial fission mediates ceramide-induced metabolic disruption in skeletal muscle. Biochem J. Dec 15 2013;456(3):427–39. doi:10.1042/BJ20130807

33. Ngo J, Choi DW, Stanley IA, et al. Mitochondrial morphology controls fatty acid utilization by changing CPT1 sensitivity to malonyl-CoA. EMBO J. Jun 1 2023;42(11):e111901. doi:10.15252/embj.2022111901

34. Buck MD, O’Sullivan D, Klein Geltink RI, et al. Mitochondrial Dynamics Controls T Cell Fate through Metabolic Programming. Cell. Jun 30 2016;166(1):63–76. doi:10.1016/j.cell.2016.05.035

35. Lee H, Yoon Y. Mitochondrial fission: regulation and ER connection. Mol Cells. Feb 2014;37(2):89–94. doi:10.14348/molcells.2014.2329

36. Adaniya SM, J OU, Cypress MW, Kusakari Y, Jhun BS. Posttranslational modifications of mitochondrial fission and fusion proteins in cardiac physiology and pathophysiology. Am J Physiol Cell Physiol. May 1 2019;316(5):C583–C604. doi:10.1152/ajpcell.00523.2018

37. Twig G, Elorza A, Molina AJ, et al. Fission and selective fusion govern mitochondrial segregation and elimination by autophagy. EMBO J. Jan 23 2008;27(2):433–46. doi:10.1038/sj.emboj.7601963

38. Ruhmkorf A, Harbauer AB. Role of Mitochondria-ER Contact Sites in Mitophagy. Biomolecules. Jul 31 2023;13(8)doi:10.3390/biom13081198

39. Krols M, van Isterdael G, Asselbergh B, et al. Mitochondria-associated membranes as hubs for neurodegeneration. Acta Neuropathol. Apr 2016;131(4):505–23. doi:10.1007/s00401-015-1528-7

40. Sakai S, Watanabe S, Komine O, Sobue A, Yamanaka K. Novel reporters of mitochondria-associated membranes (MAM), MAMtrackers, demonstrate MAM disruption as a common pathological feature in amyotrophic lateral sclerosis. FASEB J. Jul 2021;35(7):e21688. doi:10.1096/fj.202100137R

41. Giacomello M, Pellegrini L. The coming of age of the mitochondria-ER contact: a matter of thickness. Cell Death Differ. Sep 1 2016;23(9):1417–27. doi:10.1038/cdd.2016.52

42. Liu Y, Mao ZH, Huang J, et al. Mitochondria-Associated Endoplasmic Reticulum Membranes in Human Health and Diseases. MedComm (2020). Jul 2025;6(7):e70259. doi:10.1002/mco2.70259

43. Mengeste AM, Rustan AC, Lund J. Skeletal muscle energy metabolism in obesity. Obesity (Silver Spring). Oct 2021;29(10):1582–1595. doi:10.1002/oby.23227

44. Stanley WC, Lopaschuk GD, Hall JL, McCormack JG. Regulation of myocardial carbohydrate metabolism under normal and ischaemic conditions. Potential for pharmacological interventions. Cardiovasc Res. Feb 1997;33(2):243–57. doi:10.1016/s0008-6363(96)00245-3

45. Koves TR, Ussher JR, Noland RC, et al. Mitochondrial overload and incomplete fatty acid oxidation contribute to skeletal muscle insulin resistance. Cell Metab. Jan 2008;7(1):45–56. doi:10.1016/j.cmet.2007.10.013

46. Kostyo JL. A study of the glycogen levels of the rat uterus and certain skeletal muscles during pregnancy. Endocrinology. Jan 1957;60(1):33–7. doi:10.1210/endo-60-1-33

47. Steingrimsdottir T, Ronquist G, Ulmsten U. Energy economy in the pregnant human uterus at term: studies on arteriovenous differences in metabolites of carbohydrate, fat and nucleotides. Eur J Obstet Gynecol Reprod Biol. Oct 29 1993;51(3):209–15. doi:10.1016/0028-2243(93)90037-d

48. Steingrímsdóttir T, Ronquist G, Ulmsten U, Waldenström A. Different energy metabolite pattern between uterine smooth muscle and striated rectus muscle in term pregnant women. Eur J Obstet Gynecol Reprod Biol. Oct 1995;62(2):241–5. doi:10.1016/0301-2115(95)02195-d

49. Steingrimsdottir T, Ericsson A, Franck A, Waldenstrom A, Ulmsten U, Ronquist G. Human uterine smooth muscle exhibits a very low phosphocreatine/ATP ratio as assessed by in vitro and in vivo measurements. Eur J Clin Invest. Sep 1997;27(9):743–9. doi:10.1046/j.1365-2362.1997.1810726.x

50. Randle PJ, Garland PB, Hales CN, Newsholme EA. The glucose fatty-acid cycle. Its role in insulin sensitivity and the metabolic disturbances of diabetes mellitus. Lancet. Apr 13 1963;1(7285):785–9. doi:10.1016/s0140-6736(63)91500-9

51. Hue L, Taegtmeyer H. The Randle cycle revisited: a new head for an old hat. Am J Physiol Endocrinol Metab. Sep 2009;297(3):E578–91. doi:10.1152/ajpendo.00093.2009

52. Sugden MC, Holness MJ. Mechanisms underlying regulation of the expression and activities of the mammalian pyruvate dehydrogenase kinases. Arch Physiol Biochem. Jul 2006;112(3):139–49. doi:10.1080/13813450600935263

53. Murata T, Prifti KK, Ma X, Kent LN, Frolova AI, England SK. Myometrial smooth muscle Katp channel activity is attenuated in a mouse model of diet-induced obesity during pregnancy. Biol Reprod. Jun 13 2025;doi:10.1093/biolre/ioaf130

54. Doenst T, Nguyen TD, Abel ED. Cardiac metabolism in heart failure: implications beyond ATP production. Circ Res. Aug 30 2013;113(6):709–24. doi:10.1161/CIRCRESAHA.113.300376

55. Liesa M, Shirihai OS. Mitochondrial dynamics in the regulation of nutrient utilization and energy expenditure. Cell Metab. Apr 02 2013;17(4):491–506. doi:10.1016/j.cmet.2013.03.002

56. Kelley DE, He J, Menshikova EV, Ritov VB. Dysfunction of mitochondria in human skeletal muscle in type 2 diabetes. Diabetes. Oct 2002;51(10):2944–50. doi:10.2337/diabetes.51.10.2944

57. Bonnard C, Durand A, Peyrol S, et al. Mitochondrial dysfunction results from oxidative stress in the skeletal muscle of diet-induced insulin-resistant mice. J Clin Invest. Feb 2008;118(2):789–800. doi:10.1172/JCI32601

58. Wredenberg A, Freyer C, Sandstrom ME, et al. Respiratory chain dysfunction in skeletal muscle does not cause insulin resistance. Biochem Biophys Res Commun. Nov 10 2006;350(1):202–7. doi:10.1016/j.bbrc.2006.09.029

59. Pospisilik JA, Knauf C, Joza N, et al. Targeted deletion of AIF decreases mitochondrial oxidative phosphorylation and protects from obesity and diabetes. Cell. Nov 2 2007;131(3):476–91. doi:10.1016/j.cell.2007.08.047

