## Supplemental for "Maternal obesity disrupts glucose oxidation and mitochondrial fission in the myometrium at term gestation"

**Supplementary Figure S1**

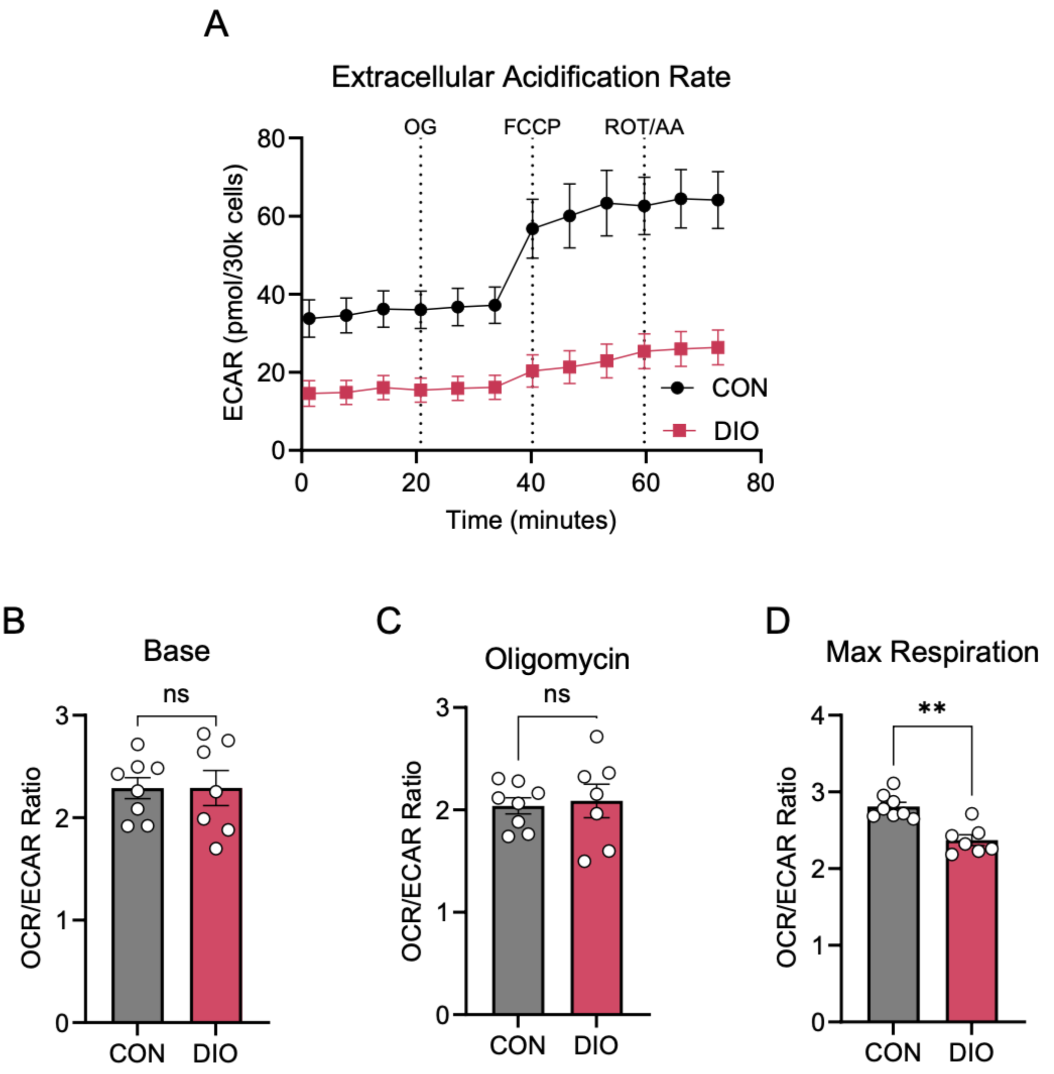


**Supplementary Figure S1**

**Oxygen consumption rate / extracellular acidification rate (OCR/ECAR) ratio is reduced at maximal respiration in DIO myometrial cells.** (A) Combined Mitochondrial Stress Test traces of ECAR for CON (n=8, grey) and DIO murine myometrial smooth muscle cells (n=7, pink). Quantification of (B) OCR/ECAR at baseline, (C) OCR/ECAR after oligomycin (OG), and (D) OCR/ECAR at maximal respiration. Data are presented as means± SEM. ***P*< 0.01 by Student *t* test. AA, antimycin A; FCCP, trifluoromethoxy carbonyl cyanide phenylhydrazone; ROT, rotenone. ns, not significant.

**Supplementary Figure S2**

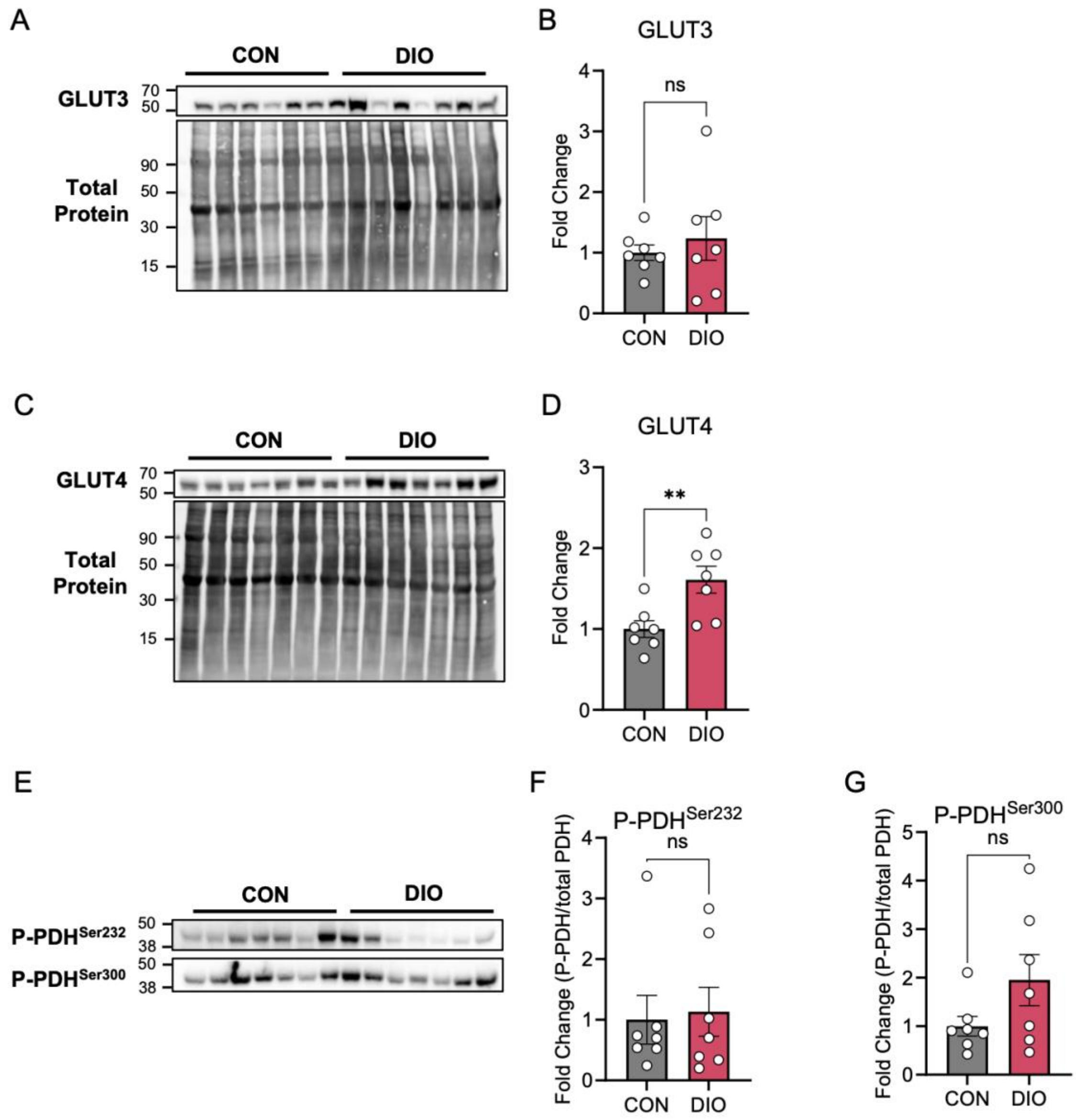


**Supplementary Figure S2**

**Pyruvate dehydrogenase phosphorylation is similar in DIO and CON myometrium.** (A) Representative Western blot probed with antibody to facilitative glucose transporter 3 (GLUT3), with corresponding total protein blot. (B) Quantification of GLUT3 relative to total protein. (C) Representative Western blot probed with antibody to facilitative glucose transporter 4 (GLUT4), with corresponding total protein blot. (D) Quantification of GLUT4 relative to total protein. (E) Representative Western blots probed with antibodies to serine 232 and serine 300 phosphorylated pyruvate dehydrogenase (P-PDH$^{Ser232}$ and P-PDH$^{Ser300}$). Quantification of (F) phosphorylated-PDH$^{Ser232}$ and (G) P-PDH$^{Ser300}$ relative to total PDH (Figure 4J). Data are presented as mean ± SEM. ***P*< 0.01 by Student *t* test. ns, not significant.

**Supplementary Figure S3**

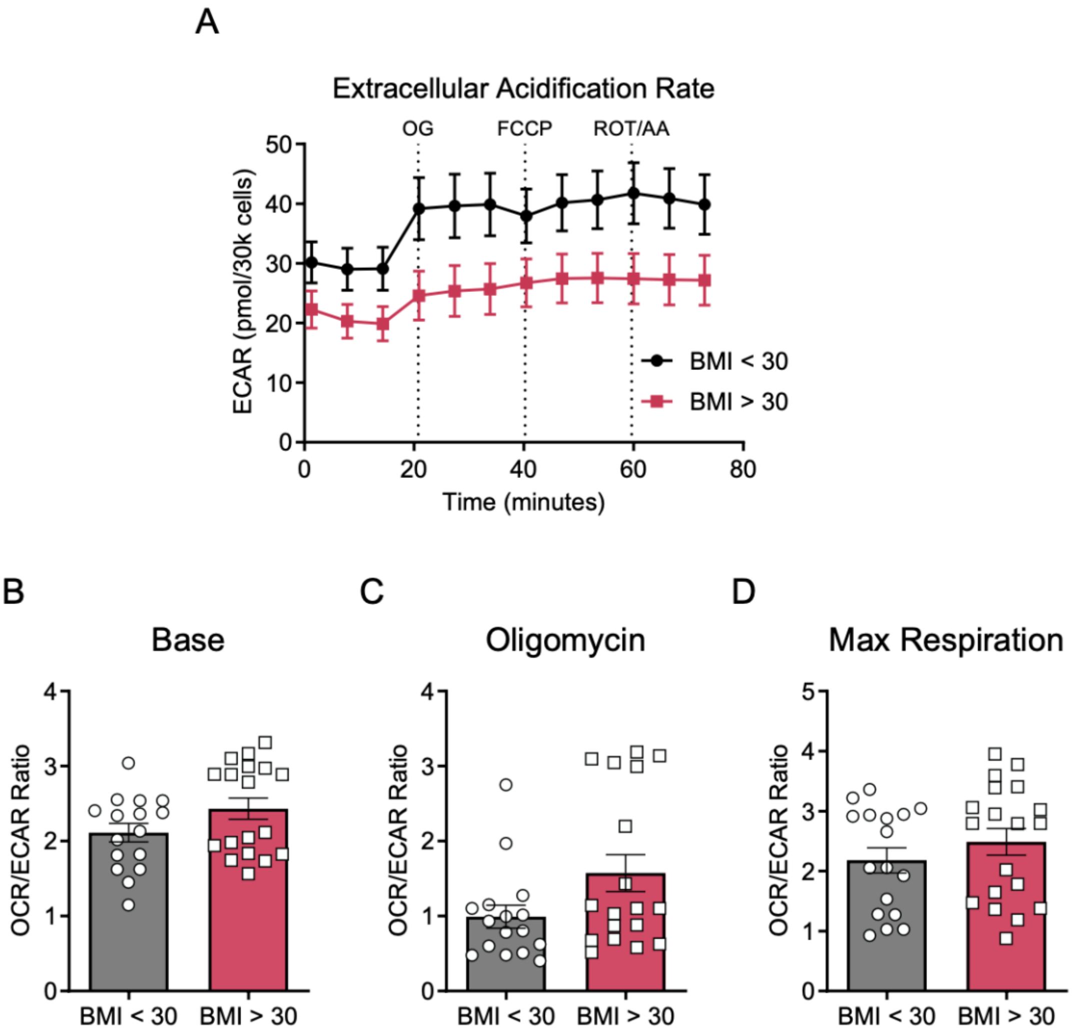


**Supplementary Figure S3**

**Body mass index is not associated oxygen consumption rate / extracellular acidification rate (OCR/ECAR) ratio in human myometrial cells.** (A) Combined Mitochondrial Stress Test traces of ECAR for human primary myometrial muscle cells from patients with body mass index (BMI) < 30 kg/m$^2$ or BMI ≥ 30 kg/m$^2$. Quantification of (B) OCR/ECAR at baseline, (C) OCR/ECAR after oligomycin (OG), and (D) OCR/ECAR at maximal respiration. Data are presented as mean ± SEM. Statistical analysis by Student *t* test. AA, antimycin A; FCCP, trifluoromethoxy carbonyl cyanide phenylhydrazone; ROT, rotenone.

**Table 1**

Table 1: Demographics

| Population characteristics | Low BMI (n=16) | High BMI (n=18) | P-value |
|---|---|---|---|
| Maternal age (y) | 29.8 +/- 5.9 | 32.2 +/- 4.4 | 0.19 |
| AMA | 5 (31.2) | 4 (26.7) | 0.70 |
| BMI, delivery | 26.2 +/- 2.4 | 48.4 +/- 11.5 | <0.0001 |
| Nulliparous | 3 (21.4) | 3 (16.7) | 1.00 |
| Parity | 1.4 +/- 1.2 | 1.8 +/- 1.9 | 0.51 |
| Gravida | 3.2 +/- 2.4 | 3.9 +/- 2.8 | 0.49 |

Values are mean +/- SD, median (IQR) or n (%)
AMA, advanced maternal age (≥35 years old); BMI, body mass index.

**Table 1**

**Patient Demographics.**
